# Programming of Embryonic Blood Brain Barrier and Neurovascular Transcriptome by an Anticipatory Acoustic Signal of Heat in the Zebra Finch

**DOI:** 10.64898/2026.01.23.701307

**Authors:** Prakrit Subba, Mylene M. Mariette, Katerina A. Palios, Michael G. Emmerson, Elisabetta Versace, Katherine L. Buchanan, David F. Clayton, Julia M. George

## Abstract

Organisms have evolved mechanisms to adjust to rapid environmental change. A dramatic example is the Australian zebra finch, where incubating parents produce an acoustic signal ("heat call") during extreme heat, triggering adaptive phenotypic plasticity in their offspring growth, thermoregulation, and reproductive success. To elucidate for the first time the molecular mechanisms underlying heat call-induced programming, we hypothesized a prenatal shift in hypothalamic gene expression, given the hypothalamus’s central role in neuroendocrine signaling controlling metabolism and growth. In addition, we tested whether prenatal heat-call exposure induces local changes in the brain, to protect this highly heat-sensitive organ from upcoming heat challenges. We exposed zebra finch embryos to chronic playback of parental heat calls or control calls, then isolated the hypothalamus for RNA sequencing to identify differentially expressed genes and gene regulatory networks. Heat-call exposure elicited modest neuroendocrine gene expression changes, but robust downregulation of genes tied to muscle contraction and cytoskeletal dynamics, with evidence of isoform usage shifts. These changes were prominently localized to hypothalamic neurovascular endothelial, mural, and ependymal cell populations, forming the blood–brain barrier (BBB). Because embryos experienced heat-associated sound, but not heat itself, and changes matched the loosening of the BBB to avoid breakages, these transcriptomic shifts likely represent an anticipatory response to enhance subsequent brain resilience to heat. Our study provides the first genome-wide characterization of embryonic hypothalamic gene expression in a songbird and reveals that prenatal acoustic cues can developmentally program neurovascular systems, expanding current understanding of developmental plasticity under climate change.

## 1 Introduction

Prenatal experiences are essential determinants of development, triggering a wide variety of changes in individual phenotype and fitness across taxa (Mariette et al., 2021; Monaghan, 2008). This process by which environmental conditions during critical early-life periods induce changes in developmental trajectories and in the resulting long-term phenotypes is called development programming (Barker, 1990; Gluckman, 2005; Monaghan, 2008; Nettle & Bateson, 2015). Biochemical cues such as maternal nutrition and hormones are well-studied drivers of development programming (Bernardo, 1996; Groothuis & Schwabl, 2008; Mariette et al., 2021; Monaghan, 2008; Mousseau & Fox, 1998). However, for other developmental drivers, such as acoustic signals that were only recently discovered (Mariette and Buchanan 2016), the underlying mechanistic pathways remain incompletely resolved (Mariette, 2024). For example, in the zebra finch (*Taeniopygia guttata*), parental vocalizations may provide embryos with a developmental cue of the immediate environmental conditions during late incubation, forecasting those upon hatching (Mariette, 2020). Adult zebra finches produce heat calls when exposed to high ambient temperatures, and more often during late incubation (Mariette & Buchanan, 2016, 2019; McDiarmid et al., 2018). Late-stage embryos exposed to these heat calls prenatally display subsequent adaptive changes postnatally on the medium- to long-term such as (i) altered growth rates, begging call production and mitochondrial functions as nestlings, and (ii) altered nest thermal preference, higher heat tolerance, and higher reproductive success as adults (Mariette & Buchanan, 2016; Pessato et al., 2022; Udino et al., 2021; Udino & Mariette, 2022). Thus, prenatal acoustic cues can produce developmental phenotypic changes in the behavior, morphology and physiology of the offspring that can be adaptive in nature to suit the environmental conditions through developmental programming (Mariette et al., 2021). However, the overarching mechanism of developmental programming in response to heat call exposure remains unknown.

Here we reasoned that postnatal phenotypic changes induced by prenatal heat call exposure would be preceded by changes in gene expression prenatally, upon receiving the acoustic cue. In particular, we hypothesized that changes in growth and thermal tolerance likely involve gene expression changes mediated by the hypothalamus. The hypothalamus is an essential neuroendocrine regulator, and neuroendocrine signaling via the hypothalamic–pituitary axes modulates processes including behavior, metabolism, thermoregulation, and growth (Debonne et al., 2008; Groothuis, 2019; Sheng et al., 2020). There is extensive evidence across taxa, including mammals, that early-life stressors can reprogram the hypothalamic–pituitary–adrenal (HPA) axis, with lasting consequences for glucocorticoid physiology and downstream developmental trajectories (Plotsky & Meaney, 1993; Welberg & Seckl, 2001). Altered HPA sensitivity occurs not only in the context of early-life stress, but also in birds, in which maternal hormones and prenatal sensory signals can modulate developmental outcomes via the HPA axis (Kraft et al., 2021; Ruiz-Raya et al., 2023). For instance, in yellow-legged gulls (*Larus michahellis*), embryos perceive prenatal cues such as adult alarm calls and shifts in light exposure and respond with coordinated changes in corticosterone levels and glucocorticoid receptor expression (Noguera & Velando, 2019; Ruiz-Raya et al., 2023). However, evidence linking such HPA-axis modifications to changes in hypothalamic gene regulation itself is still limited (Avishai-Eliner et al., 2001; Korosi et al., 2010; Marasco et al., 2016; Yan et al., 2020; Zimmer & Spencer, 2014). We therefore specifically targeted the hypothalamus to test the primary hypothesis that neuroendocrine pathways control the thermoregulatory programming of individual traits observed in nestlings and in adulthood.

Beyond neuroendocrine programming, upon receiving an anticipatory signal of heat, the brain itself, one of the most heat-sensitive organs of the body (Bain et al., 2015; Yoneda et al., 2024), may also undergo local readjustments to allow it to withstand high ambient temperatures. Under hyperthermia (i.e. above normal body temperature), the mechanisms to allow brain cooling or avoid brain damage are not fully elucidated in birds and mammals, but involve changes in blood flow and in oxygen assimilation (Bain et al., 2015; Porter & Witmer, 2016). Across taxa, brain blood supply is controlled by a diverse range of non-neuronal vascular-associated cells, including endothelial cells lining the interior of blood vessels and which constitute the blood-brain barrier (BBB), supported by muscle-like mural cells (e.g. vascular smooth muscle cells, VSMC; and pericytes) and glial cells (e.g. astrocytes, microglia) (Bain et al., 2015; McConnell & Mishra, 2022; Paton et al., 2023). These non-neuronal cells are increasingly understood as being highly plastic and stress-sensitive, including to heat stress (Hayes et al., 2022; Phan et al., 2025), with heatstroke in humans and model mammals characterized by cellular damage following hemorrhage and inflammation, including in the hypothalamus (Yoneda et al., 2024). Remarkably, these cells are also subject to developmental programming (Smolders et al., 2018) with recent evidence confirming that early-life stress can have lifelong detrimental effects on the blood-brain barrier, by interfering with its development (Zhao et al., 2022). Therefore, a non-mutually exclusive hypothesis to that of neuroendocrine programming above is that heat-calls trigger preventive changes to non-neuronal cells to better protect the hypothalamus from permanent damage by heat stress.

To test whether embryonic exposure to heat calls influences embryonic hypothalamic gene expression, we exposed zebra finch eggs to playback of parental heat calls or control calls during late-stage incubation. We precisely targeted the hypothalamic region using medial forebrain punches and obtained RNA sequencing (RNA-seq) data to capture transcriptomic responses. To the best of our knowledge, to date no study has quantified embryonic hypothalamic gene expression, neither in songbirds, nor in relation to any prenatal environmental stimuli with the aim of assessing the functional programming pathways. To test for coordinated changes in gene expression associated with playback condition that would reveal the endocrine or other functional pathways underlying changes to developmental trajectories and postnatal phenotypes, we conducted both a genome-wide analysis of differentially expressed transcripts, and an analysis specifically focused on genes annotated for hypothalamic function. To test our second hypothesis of localized preparatory changes in the brain, we first used the same genome-wide analysis that systematically interrogates gene expression changes between playback groups, focusing both on individual genes and on isoform-level shifts. Next, to more comprehensively characterize transcriptional changes, we applied weighted gene co-expression network analysis (WGCNA), a network approach that clusters genes with correlated expression into modules and relates these modules to specific hypothalamic cell types and candidate regulatory hub genes. Our study brings unique insights into the impact of prenatal sound on gene regulation in the embryonic brain and reveals novel heat-adaption responses consistent with cerebral protection against heat insults.

## 2 Results

### 2.1 Targeted Hypothalamus Gene Expression Analysis

Medial hypothalamic brain punches were collected from E14 embryos (i.e. day before hatch) that had been exposed to playbacks of either heat calls or control calls for 9 hours per day from incubation day 10 (E10) to 13 (E13) (n= 10 heat call, 9 control call exposed embryos; Fig. 1). In contrast to our primary hypothesis, our genome-wide expression profiling (see below) did not identify changes in genes or processes typically associated with hypothalamic function. As an adjunct to this approach, we compiled a literature-derived list of 143 hypothalamic genes (Table S1) and performed a targeted differential expression analysis on this subset, thereby reducing the multiple-testing penalty inherent in the whole-transcriptome analysis. The expression of no particular genes reached significance at an FDR threshold of 0.05. Nonetheless, 12 genes showed differential expression at an unadjusted significance threshold (Wald test p < 0.05), although adjusted p-values exceeded 0.25 (Fig. S1).

**Fig. 1:**
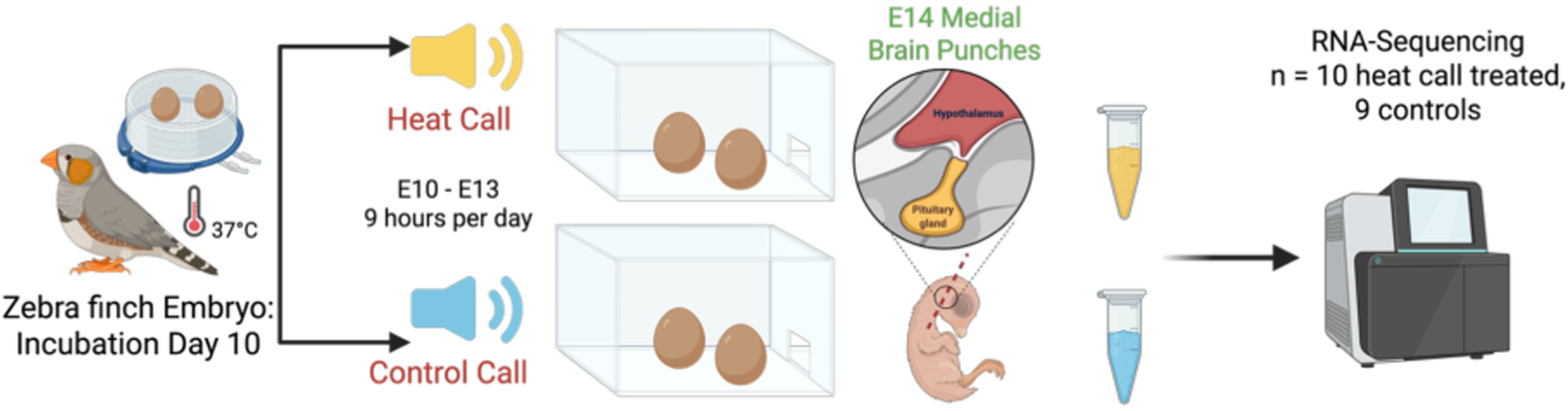
Experimental design for RNA-sequencing in late-stage zebra finch embryos. Schematic of the experimental design for RNA-sequencing in late-stage zebra finch embryos exposed to chronic heat call (n=10) or control call (n=9) playback. Created using BioRender.com

Among these candidates, the glucocorticoid receptor gene *NR3C1* showed the strongest evidence for modulation, with reduced expression in heat-call–exposed embryos relative to control call playbacks (shrunken log2 fold change = 0.26, p = 0.0045, FDR = 0.25). Aryl hydrocarbon receptor nuclear translocator gene *ARNT* was also downregulated (shrunken LFC = 0.13, p = 0.0088, FDR = 0.25). In contrast, several neuroendocrine genes showed modest increases in expression following heat-call exposure: corticotropin-releasing hormone receptor 1 gene *CRHR1* (shrunken LFC = 0.068, p = 0.013, FDR = 0.29), growth hormone-releasing hormone gene *GHRH* (shrunken LFC = 0.021, p = 0.0083, FDR = 0.25), neurotensin gene *NTS* (shrunken LFC = 0.051, p = 0.0075, FDR = 0.25), and somatostatin gene *SST* (shrunken LFC = 0.066, p = 0.027, FDR = 0.42) (Fig. S1 and Table S1).

### 2.2 Genome-wide Downregulation of Muscle and Cytoskeletal Genes in the Hypothalamus in Heat Call Exposed Embryos

Across the 19 samples, RNA-seq yielded a mean effective library size of 33.8 million gene-level counts (median 31.8 million; range 26.4–62.6 million; n = 19 individuals), and 19,137 genes with at least 10 total counts were retained for differential expression analyses. To characterize global gene expression patterns in the dataset, principal component analysis of variance-stabilized counts (19,137 genes, 19 samples) revealed pronounced transcriptomic sex differentiation, with PC1 (48% of variance) strongly separating males and females (r = 0.989; Fig. S2). To our knowledge, this represents the first demonstration of sex-differentiated gene expression in the embryonic songbird hypothalamus. Besides sexual differentiation, both PC1 and PC2 (17% of variance) captured moderate separation by playback condition (r = −0.549 and r = 0.477, respectively), with only weak association with sex for PC2 (r = 0.117; Fig. S2). Therefore, beyond sex differences, prenatal acoustic experience explained a noticeable amount of variation in overall hypothalamic gene expression.

Using a DESeq2 model including sex and playback condition (∼ sex + playback condition), 49 genes were differentially expressed between heat call and control embryos (padj < 0.05). The response was strongly asymmetric, with 48 genes downregulated and 1 gene upregulated in heat-call–exposed embryos relative to control call playback (Fig. 2A, Table S2). A secondary model including the interaction term (∼ sex + playback condition + sex:playback condition) identified five genes with significant sex-by-treatment interactions (padj < 0.05): three sex-linked genes (*HDHD2* on Z chromosome; *LOC116806857* and *LOC116806879*/*KCMF1* on W chromosome) and two autosomal genes (*ABRACL* and *PTPRM*), indicating that some transcriptional responses to heat-call exposure differ between males and females.

**Fig. 2:**
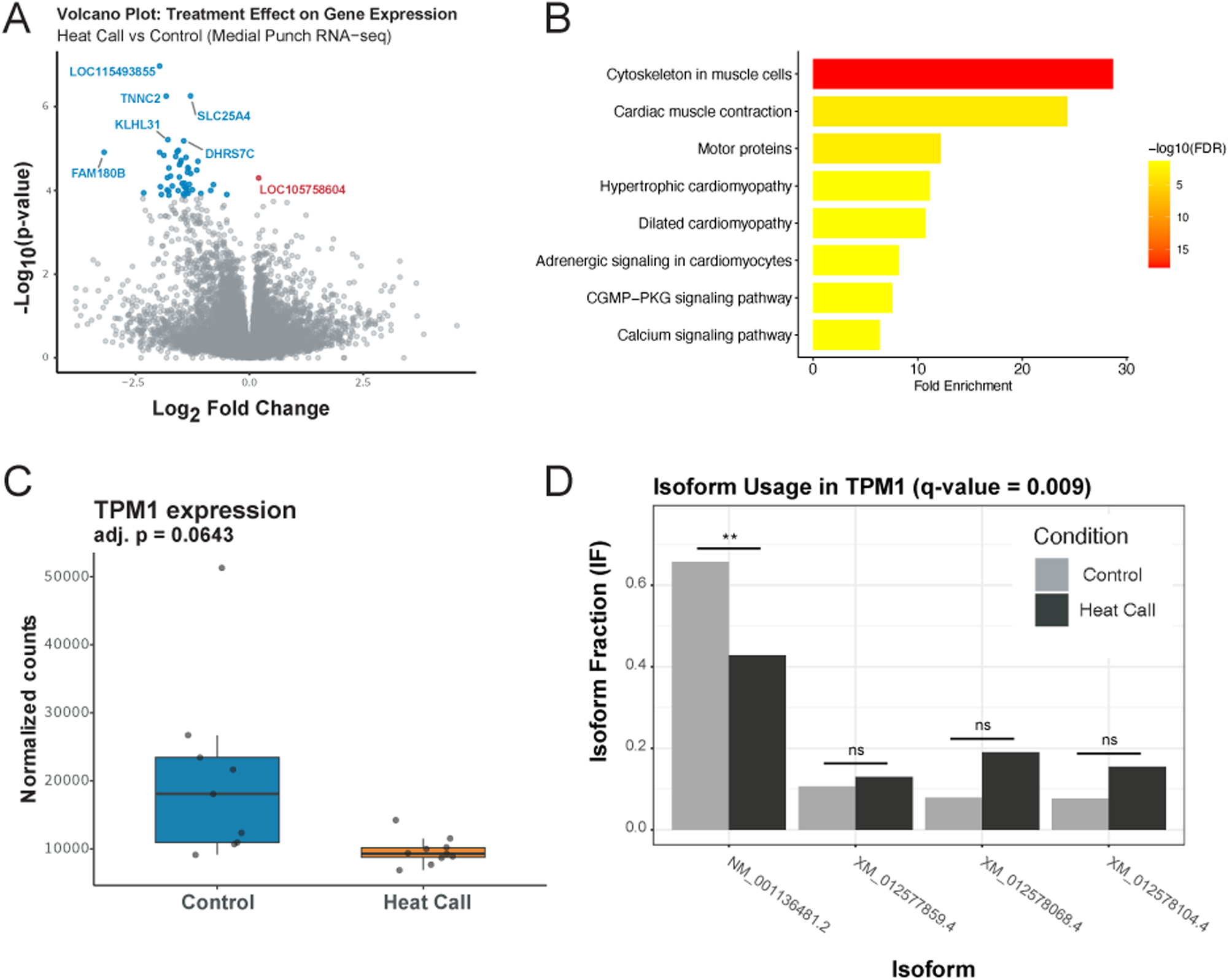
RNA-seq analysis of medial hypothalamic punches from late-stage zebra finch embryos exposed to chronic heat call or control call playback. Medial hypothalamic punches were collected from n = 10 heat call and n = 9 control embryos (one punch per embryo; 19 biological replicates). (A) Volcano plot showing log2 fold change (x-axis) and −log10(p-value) (y-axis) for individual genes; blue and red points indicate genes with Benjamini–Hochberg adjusted p-value (padj) < 0.05 from DESeq2 likelihood ratio tests (blue = downregulated, red = upregulated; 49 DEGs total: 48 downregulated, 1 upregulated). (B) Pathway enrichment of treatment-responsive DEGs analyzed in ShinyGO (v0.85.1, KEGG database) using hypergeometric tests with Benjamini–Hochberg FDR correction; bars show fold enrichment, color encodes −log10(FDR). (C–D) Nominal downregulation of TPM1 (DESeq2 p_adj_=0.064) and differential isoform usage; panel D shows isoform fractions (y axis) for individual TPM1 transcripts (x-axis) in control (grey) and heat call (black) conditions, with significant differential usage from IsoformSwitchAnalyzeR DEXSeq tests (two-sided FDR-corrected; **q < 0.01, |ΔIF| ≥ 0.05).

Pathway enrichment analysis of the 48 downregulated DEGs revealed significant over-representation of muscle cytoskeletal and excitation–contraction pathways (enrichment FDR < 0.05; Fig. 2B). The top enriched terms were Cytoskeleton in muscle cells (FDR < 0.01; fold enrichment = 28.69), Cardiac muscle contraction (FDR < 0.01; fold enrichment = 24.30), and Motor proteins (FDR < 0.01; fold enrichment = 12.20). These pathway-level signals mirror the downregulation of multiple canonical contractile/cytoskeletal genes in the dataset, including *TPM1*/*TPM2*/*TPM4*, *ACTC1*/*ACTA1*, *MYL2*/*MYL3*, *TNNC1*/*TNNT2*, *DES*, and *LMNA* (Table S3).

Together, these results support the conclusion that a focused, muscle-like transcriptional program, presumably related to blood-flow control by muscle contraction and cytoskeleton function of mural cells (Paton et al., 2023) is selectively repressed in the medial hypothalamus region following prenatal heat-call exposure.

### 2.3 Heat Call-Induced Isoform Switching and Alternative Splicing

Even with equal level of expression, genes may differ in which isoform is produced between playback groups. Therefore, to investigate transcript isoform changes associated with developmental programming in heat call exposed embryos, we conducted an isoform switch analysis using the transcriptome-wide RNA-seq data. After filtering to remove 20,682 low-expression and single-isoform gene transcripts, we investigated differential isoform usage in 23,054 of the remaining isoforms. A total of 26 genes showed significant treatment-specific isoform switching between playback treatments (FDR < 0.05; differential isoform fraction ≥ 0.05) (Table S4). Overall, exon skipping was the dominant alternative splicing event, accounting for 17 events among the 26 switching isoforms (Table S5). Within these, most genes showed decreased exon skipping (i.e., increased exon inclusion) in heat call-treated embryos, with the enrichment analysis showing a nominally significant difference (proportion of gains = 0.24, p = 0.049, q = 0.34; total ES events = 17) (Table S5). Other alternative splicing events (alternative splice sites, transcription start/stop sites, mutually exclusive exons, and intron retention) were less common and did not show significant differences between treatment (all q ≥ 0.77) (Table S5).

Among the significant switches, three genes showed especially strong differential isoform usage between playback groups: *TLK2,* involved in DNA repair and chromatin maintenance (q < 0.001, ΔIF = −0.06), *LOC100228270* (q = 0.009, ΔIF = −0.26), and *TPM1,* encoding Tropomyosin 1 involved in cytoskeleton and muscle contraction, and whose excess has been linked to neuroinflammation by microglia activation (q = 0.009, ΔIF = −0.23; Fig. 2D, Table S4) (R. Li, Liang, et al., 2022; R. Li, Zhang, et al., 2022). All three genes displayed a lower fraction of the most common isoform in heat-call embryos compared to control call playback. Interestingly, in the genome-wide analysis, *TPM1* also showed a borderline gene-level downregulation (log2 fold change = −1.11, padj = 0.064, baseMean = 14769.90; Fig. 2C). This result provides convergent evidence for reduced *TPM1* activity, both in overall expression and in isoform usage, following prenatal heat-call exposure. Overall, these results show that heat-call exposure predominantly led to increased exon inclusion, while most other alternative splicing modes remained unchanged.

### 2.4 Heat Call Treated Embryos Display Coordinated Downregulation of Actin Cytoskeletal and Muscle Development Genes in Neurovascular Cells

To assess possible functional changes, we identified coordinated changes in hypothalamic gene expression at the level of biological pathways by conducting a WGCNA analysis. We used DESeq2 variance-stabilized expression values for 21,407 genes detected in the medial hypothalamus RNA-seq dataset across 19 samples. All genes detected by the gene-level quantification step were retained for co-expression network construction, without additional variance-based pre-filtering, and module detection was performed on this full gene set. We identified 28 distinct modules, which each contains a group of highly correlated genes with similar expression patterns across samples, with sizes ranging from 76 to 5,629 genes in each module (Fig. 3). In our module-trait correlation analysis, the green module (n = 463 genes) showed the strongest negative correlation with heat call response (r = -0.60, p < 0.01) (Fig. 3). Further, the brown (r = -0.46, p = 0.047) and red (r = -0.46, p = 0.045) modules were also negatively correlated with heat call playbacks (Fig. 3). The red module also involved genes related to muscle contraction and development (Table S3). The brown module (n = 1,363 genes) showed the strongest correlation with sex, indicating upregulation in females (r = 0.98, p < 0.001) (Fig. 3). No other modules had significant module-trait correlations.

**Fig. 3:**
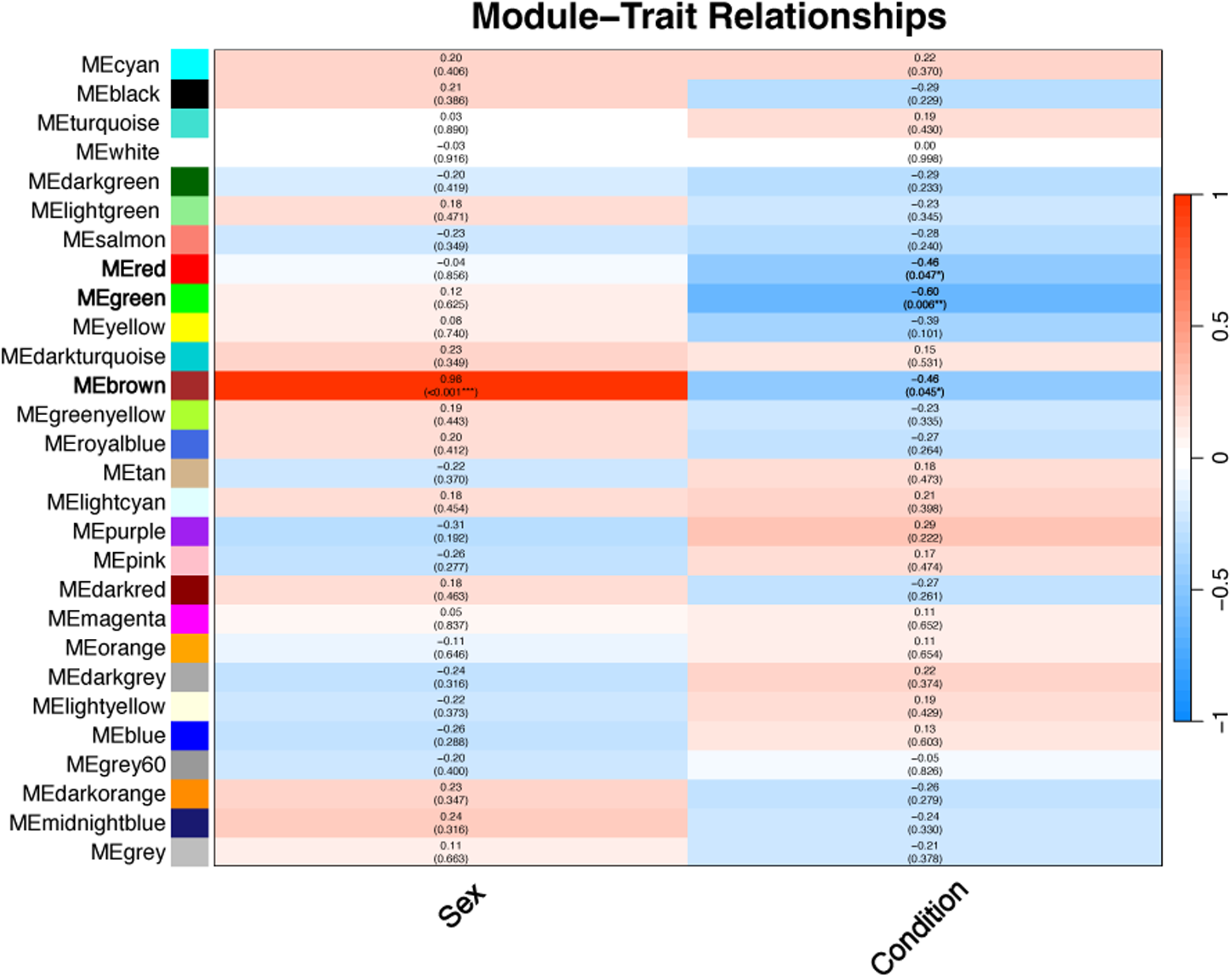
Module–trait relationships between WGCNA eigengenes and sex or playback condition. Heatmap showing Pearson correlation coefficients (cell values) and two-sided Student’s t-test p-values (parentheses) between WGCNA module eigengenes and sex (left) or playback condition (right), derived from medial hypothalamic RNA-seq (n=19: 10 heat call, 9 control embryos), with males and control call used as the reference respectively. Color intensity reflects association strength and direction (red = positive; blue = negative).

Next, we performed a gene ontology (GO) term analysis to investigate the biological functions associated with each of these modules correlated with heat call playback. We found that the green module was significantly enriched in terms such as Myofibril assembly (GO:0030239), Cytoskeleton in muscle cells (Path:tgu04820), Muscle structure development (GO:0061061), and Contractile muscle fiber (GO:0043292) (FDR < 0.01) suggesting the module’s involvement in muscle developmental and cytoskeleton-related programs (Fig. 4B). *TPM1*, a gene marginally differentially expressed and differentially spliced in heat call exposed embryos (see above), was also a member of the green module enriched muscle programs (Fig. 4A, Table S3). Intramodular connectivity analysis in the green module identified a tightly connected network centered on five hub genes with the highest intramodular connectivity (kWithin). These hub genes are typically involved in the functioning of cardiac and skeletal muscle cells, but can also be expressed to a lesser extent in the brain, with functions thought to be related to neuronal development and cytoskeletal structure (Blech-Hermoni et al., 2023; Nelson et al., 2021), such as ADP-ribosylhydrolase-like 1 (*ADPRHL1*), Cardiomyopathy Associated 5 (*CMYA5*), LIM Domain Binding 3 (*LDB3*), Kelch-like Family Member 40 (*KLHL40*), and Titin (*TTN*) and have strong interactions (adjacency > 0.4) with associated neighbor genes in the green module (Fig. 4B).

**Fig. 4:**
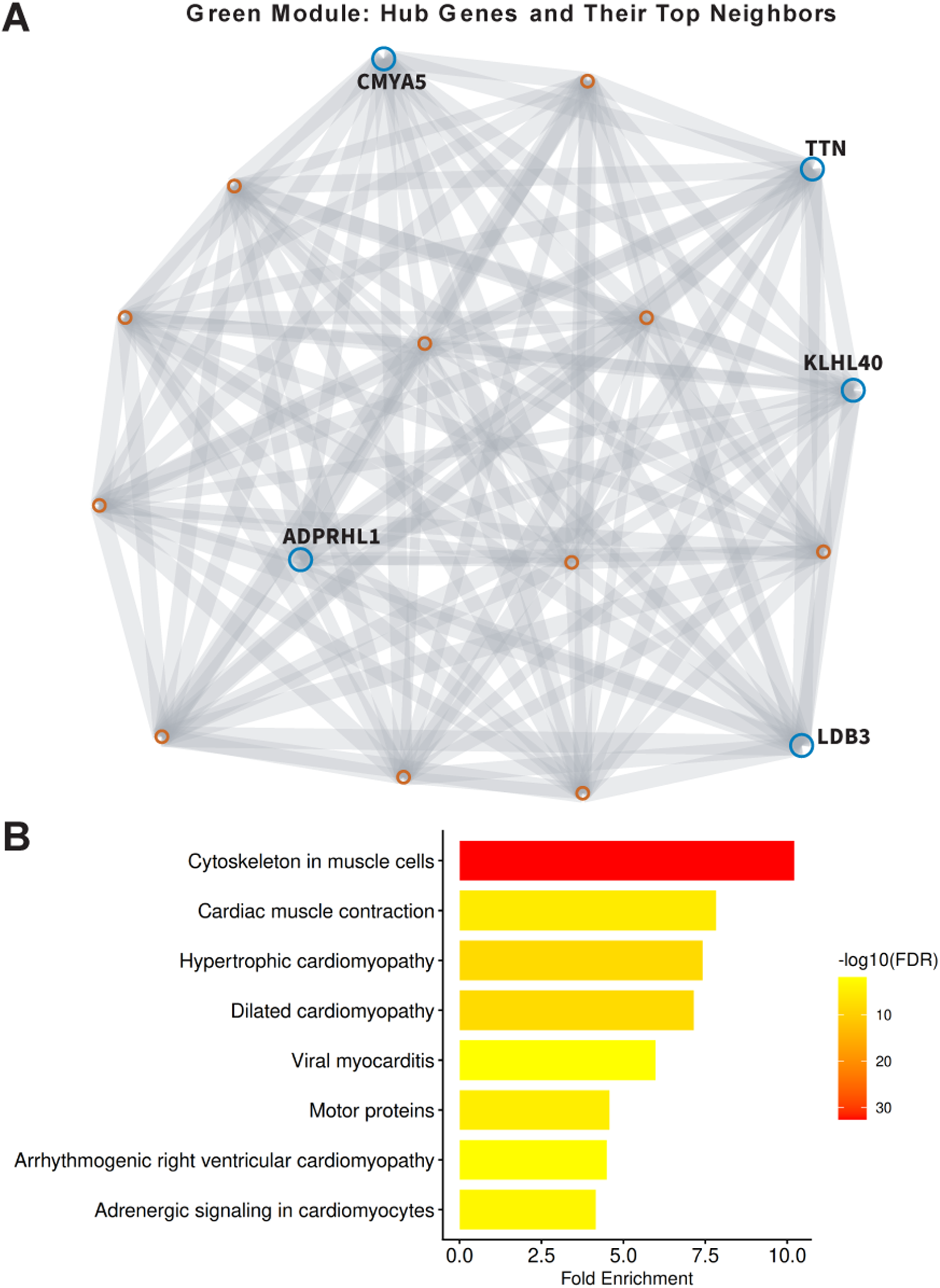
Green WGCNA module structure and functional enrichment. (A) Green module co-expression network from WGCNA of hypothalamic RNA-seq data (n=19 embryos: 10 heat call, 9 control) showing top 5 hub genes (larger blue circles; ranked by intramodular connectivity kWithin) and their strongest co-expressed neighbors (smaller red circles; adjacency > 0.4). Edge thickness indicates pairwise co-expression strength. (B) Pathway enrichment analysis for green module genes analyzed in ShinyGO (v0.85.1, KEGG database) using hypergeometric tests with Benjamini–Hochberg FDR correction; bars show fold enrichment, color encodes −log10(FDR).

To identify the specific hypothalamic cell types in which gene expression changes occurred, we conducted cell-type enrichment analysis within each module (see Methods). The results showed that the green module genes were significantly enriched in mural cells (i.e., vascular smooth muscle cells VSMC and pericytes; FDR < 0.001), endothelial cells (FDR = 0.015), and ependymal cells (FDR = 0.015; Fig. 5). Genes in the brown and red modules showed no significant cell-type enrichment for hypothalamic cell types.

**Fig. 5:**
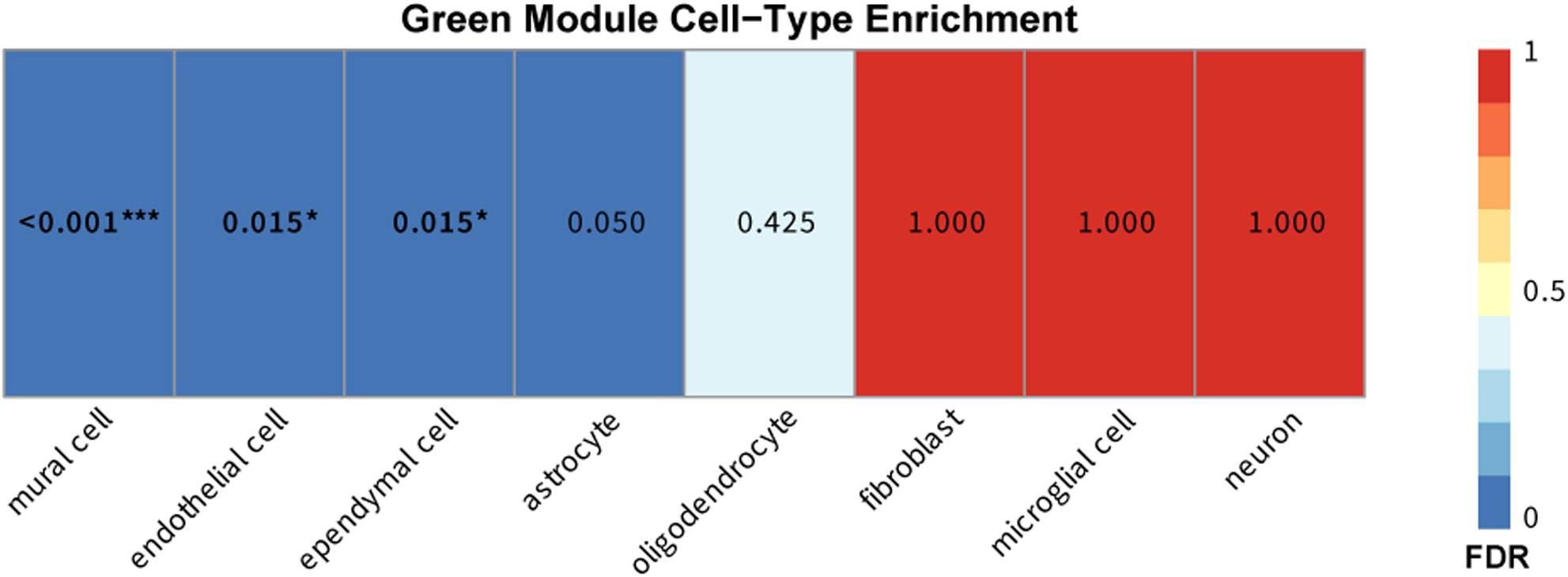
Cell-type enrichment of the green WGCNA module using Hypomap markers. Heatmap of FDR-corrected p-values from one-sided hypergeometric tests assessing overrepresentation of green WGCNA module genes (from n=19 hypothalamic samples: 10 heat call, 9 control; converted to human homologs via biomaRt) within Hypomap cell-type marker gene sets. Benjamini–Hochberg FDR correction applied across all module-by-cell-type tests. Dark blue indicates strong enrichment; *FDR < 0.05, **FDR < 0.01, ***FDR < 0.001.

### 2.5 Digital Deconvolution of Acoustic Playback Exposed Embryos Shows Enrichment of Neuronal and Glial Cell Types

To estimate the cellular composition of bulk hypothalamic tissue across all embryos, we employed CIBERSORTx deconvolution and derived cell-type-specific relative fractions representing within-sample RNA contributions. Oligodendrocytes (median: 0.403 [IQR: 0.017]), neurons (median: 0.374 [IQR: 0.061]), astrocytes (median: 0.124 [IQR: 0.018]), and microglia (median: 0.071 [IQR: 0.010]) dominated the cellular milieu, collectively representing over 95% of the estimated RNA contribution (Fig. S4). Minor populations included mural cells (median: 0.014 [IQR: 0.015]), ependymal cells (median: 0.007 [IQR: 0.011]), and endothelial cells (median: 0.005 [IQR: 0.004]), with fibroblasts near absent (median: 0.000; mean: 0.001) (Fig. S4).

Across all samples, the CIBERSORTx deconvolution met stringent diagnostic criteria (goodness-of-fit p < 0.0001, Pearson r > 0.89, RMSE 0.70–0.75; Table S6). This cellular composition is consistent with parenchyma-enriched hypothalamic punches, confirming that the dissected tissue primarily captured neural and glial cell types rather than surrounding non-neural tissues. The high proportion of neurons and glia relative to vascular-associated cells (endothelial and mural cells comprising <2% of RNA contribution) indicates that transcriptional effects localized to vascular compartments would be substantially diluted in bulk RNA-seq profiles. Despite their minimal RNA contribution (<2%), vascular-associated cells exhibited detectable transcriptional responses to playback exposure (see Section 2.4), consistent with large-magnitude gene expression changes within these cell populations that are sufficient to be detected in bulk profiles.

## 3 Discussion

In recent years, sound has emerged as a novel source of information for adaptive developmental programming in embryos (Mariette & Buchanan, 2016; Noguera & Velando, 2019; Ruiz-Raya et al., 2023). However, the impact of prenatal sound on developmental processes, and the mechanisms through which this occurs, are yet to be fully understood. In this study, we characterized embryonic hypothalamic gene expression in response to heat-call exposure to identify possible systemic and localized changes enabling a heat-adapted phenotype. To our knowledge, our study provides the first comprehensive transcriptomic characterization of the prenatal zebra finch hypothalamus. In addition, we found that prenatal heat call exposure can indeed lead to transcriptomic changes in the avian embryonic brain. Contrary to our primary hypothesis, neuroendocrine genes traditionally associated with hypothalamic function represented only a small fraction of the transcriptomic response, this signal was detected only in a targeted investigation of hypothalamic function genes to circumvent the limitations of multiple hypothesis testing and fell below the corrected significance level. Instead, the primary transcriptomic response to heat-calls involved genes associated with muscle function and cytoskeleton, localized to hypothalamic cell clusters annotated as mural cells (smooth muscle cells and pericytes), endothelial cells of the BBB, and tanycytes (ependymal cells of the third ventricle).

Most of these genes were downregulated by heat call exposure and displayed coordinated, systematic expression changes in our WGCNA analyses. These findings suggest a reprogramming of vascular regulation in the hypothalamus that may enhance blood-brain barrier integrity, a localized neuroprotective adaptation rather than a systemic orchestrating mechanism. If this neurovascular programming persists post-hatch, enhanced blood-brain barrier integrity could provide neuroprotection during heat exposure, potentially contributing to the increased heat tolerance and altered thermal preferences previously observed in heat-call-exposed offspring (Bain et al., 2015; Mariette & Buchanan, 2016; Porter & Witmer, 2016; Udino et al., 2021; Udino & Mariette, 2022).

However, such localized hypothalamic vascular changes are unlikely to account for organismal-level phenotypes documented in earlier studies, including altered growth trajectories and mitochondrial function (Mariette & Buchanan, 2016; Udino et al., 2021), indicating that additional systemic mechanisms which develop post hatch, possibly mitochondrial or other non-hormonal pathways, remain to be identified. Remarkably, embryos in our experiment were not exposed to harmful heat, but only to an acoustic signal of heat, demonstrating the potency of sensory-mediated developmental plasticity and opening unique research opportunities for understanding how prenatal acoustic cues shape physiological systems.

The BBB changes we observed involve the neurovascular unit, an integrated system composed of endothelial cells, vascular smooth muscle cells (VSMCs), astrocytes, pericytes, and neurons (McConnell & Mishra, 2022). These cells work together to control blood flow, vascular tone, nutrient delivery, waste removal, and protection from harmful substances. Endothelial cells line the interior of blood vessels by forming tight junctions that control the exchange of substances between the bloodstream and the brain (Atochin & Huang, 2011; Hogan-Cann et al., 2019; McConnell & Mishra, 2022; Stobart et al., 2013). Wrapped around brain blood vessels, mural cells (VSMCs and pericytes) contribute to blood-brain barrier integrity alongside endothelial cells and astrocytes. While the role of endothelial cells has long been investigated in medical science, current research is revealing key functionalities of mural cells and their involvement in neurovascular health and neurodegeneration (Vanlandewijck et al., 2018). Notably, VSMCs exhibit remarkable phenotypic plasticity in response to environmental cues, ranging from a "contractile" phenotype characterized by high expression of contractile proteins (smooth muscle actin, myosin heavy chain, tropomyosin) to a "synthetic" phenotype with reduced contractile gene expression and elevated proliferation, migration, and extracellular matrix production (Hayes et al., 2022; Vanlandewijck et al., 2018). Embryonic VSMCs normally exhibit a synthetic phenotype and mature to the contractile state in adulthood (Ball et al., 2010; Owens et al., 2004). Our results showing downregulation of contractile marker genes, including *TPM1*, a key regulator of VSMC contractile phenotype (S. Li et al., 2025), as well as genes associated with cytoskeleton in muscle cells and cardiac muscle contraction pathways (Green module in Section 2.4), suggest that heat-call exposure may maintain or enhance the embryonic synthetic VSMC phenotype during prenatal hypothalamic development. This retention of the synthetic state could facilitate vascular remodeling and barrier adaptation in preparation for post-hatch thermal challenges.

It is well established that, under normal temperature range, blood flow adjusts to temperature (Bain et al., 2015; Phan et al., 2025), including via temperature-sensing channels in VSMC and endothelial cells that regulate Ca^2+^ influx (Hu & Zhang, 2024; Kuppusamy et al., 2021; Phan et al., 2025). However, whether the brain possesses defense mechanisms specifically against hyperthermia is unclear, with evidence possibly trending towards the negative (Bain et al., 2015). For example, in birds, the rete ophthalmicum, a highly vascularized area behind the eye that functions as a heat exchanger (Arad & Midtgard, 1990; Bech & Midtgård, 1981), only cools a fraction of the blood reaching the brain, and proportional cooling does not increase at higher temperature (Jessen, 2001; Porter & Witmer, 2016). Likewise in humans, because of changes in respiration and blood flow for body thermoregulation, blood flow to the brain decreases at high temperatures, which reduces rather than enhances brain cooling (Bain et al., 2015). Furthermore, at the cellular level, it is not clear which changes in endothelial, mural and glial cells triggered by high temperatures may be beneficial or detrimental (Bain et al., 2015; Boorman et al., 2023; Phan et al., 2025). Under hyperthermia, heat-stress studies show a breakdown of endothelial tight junctions, mediated in part by pro-inflammatory molecules, and which leads to cerebral edema (Bain et al., 2015; Kiyatkin & Sharma, 2009; Yoneda et al., 2024). However, it has been suggested that, in the initial stages of brain challenges, the loosening of the blood brain barrier tight junctions may occur to prevent structural damage to the BBB (Huber et al., 2001). Therefore, our finding showing the downregulation of contractile and cytoskeleton function of endothelial and mural cells likely represents such adaptive modifications for BBB protection. Furthermore, because embryos were exposed to a signal of heat, but not to heat itself, it is highly unlikely that the transcriptomic changes we observed correspond to heat-induced damages, which instead typically involve an upregulation of oxidative stress, mitochondrial dysfunction and inflammation pathways (Fang et al., 2024; Yoneda et al., 2024). Investigating heat call effects therefore represents a unique opportunity for disentangling adaptive protective responses from detrimental cellular damages induced by heat-stress.

Our study also brings novel insights into the developmental programming of the BBB and neurovascular unit of the hypothalamus. The changes we observed were not an immediate response to playback (stopped 15hrs prior, the night before sampling), but a persistent programming after multiple days of acoustic exposure, potentially analogous to an acclimatization period. Indeed, heat tolerance in humans (Bain et al., 2015), and BBB resistance to heat and posterior recovery (Jeliazkova-Mecheva et al., 2006) have been shown to be improve after heat-acclimatization.

Exposure to heat calls during prenatal life may therefore provide an opportunity for triggering defense mechanisms prior to heat exposure upon hatching. Besides loosening of BBB tight junctions, heat-call exposure may induce changes to counter the heat-induced inflammation in the brain that has been linked to edema in heatstroke patients (Yoneda et al., 2024), and which could also potentially be involved in the formation of edema in zebra finch embryos suffering from hypoxia at high incubation temperature (Choi et al., 2025). At least in mammals, the inflammation response is in part triggered by *TPM1* that activates microglia (R. Li, Liang, et al., 2022; R. Li, Zhang, et al., 2022); and microglia proliferation has recently been identified as the underlying mechanism for the programming of the BBB, leading to lifelong brain inflammation following prenatal exposure to maternal immune activation (Zhao et al., 2022). Therefore, the downregulation of *TPM1* and isoform switches that we observed in zebra finch embryos exposed to heat calls, are consistent with an anti-inflammatory response in the brain. Together with those on BBB loosening, these findings provide strong support for our secondary hypothesis that heat-call exposure triggers local adjustments in the hypothalamus that are specifically directed at countering heat damage in the brain.

Contrary to our initial hypothesis, the heat-adapted phenotype triggered by prenatal heat-call exposure and observed in nestlings and adults may not primarily be driven by neuroendocrine mechanisms. Our hypothesis on endocrine programming by prenatal heat-calls received only weak support in this study, with targeted analyses of hypothalamic function genes – that in part allowed circumventing the limitations of multiple hypothesis testing – falling below corrected significance thresholds. Consistent with this finding, heat-call exposure reduced nestling heat-shock protein levels in hot conditions and their heterophile-to-lymphocyte (H/L) ratio (a marker of chronic stress), but did not alter plasma corticosterone levels (Udino et al., 2024), which argues against a neuroendocrine control. Nonetheless, postnatal hypothalamic gene regulation should be directly investigated, to test whether the slight neuroendocrine programming we detected in embryos may strengthen postnatally when phenotypic changes in growth and begging emerge in response to temperature (Mariette & Buchanan, 2016).

Alternatively, non-hormonal pathways may instead orchestrate the postnatal heat-adapted phenotype. Changes in nestling mitochondrial function following heat-call exposure point to metabolic reprogramming as one such pathway (Udino et al., 2021). Additionally, in adult zebra finches, acute heat challenge upregulated the dopaminergic pathway in the telencephalon, with gene expression levels positively correlated to individual panting behavior (Lipshutz et al., 2022). Since panting in adulthood varies with early-life exposure to heat-calls (Udino & Mariette, 2022), dopaminergic programming may also play a role in shaping the heat-call induced phenotype. Lastly, the localized vascular changes to the BBB and mural cells we observed could also in part contribute to some traits of the postnatal heat-adapted phenotype if they persist and reduce brain susceptibility to heat damage on the long-term. Namely, they could contribute to the higher heat tolerance observed in heat-call-exposed birds at adulthood, which was not explained by more efficient evaporative cooling (Pessato et al., 2022), as well as their preference for hotter microsites (Mariette & Buchanan, 2016).

One limitation is that embryonic gene expression was measured at a single post-exposure timepoint (E14), and follow-up sampling across postnatal stages will be required to determine whether neurovascular transcriptional changes persist or evolve after hatching. Although our deconvolution analysis confirmed that tissue punches were parenchyma-enriched and identified the primary cellular targets of heat-call programming, future integration of spatial transcriptomics with targeted histology would provide finer anatomical resolution of responsive transcripts within hypothalamic subregions. Another consideration is sex: we observed pronounced transcriptomic sex differentiation in the embryonic hypothalamus (PC1: 48% variance; Fig. S2), indicating that sex is a major axis of variation at this developmental stage. Importantly, although sex ratios were imbalanced between playback groups, multiple lines of evidence argue against sex confounding the main playback conclusions. First, the modules most strongly associated with playback (green and red) showed no detectable association with sex (green: r = 0.12, p = 0.625; red: r = -0.04, p = 0.856; Fig. 3), indicating qualitatively distinct axes of variation for sex versus playback. Second, the module most strongly associated with sex (brown; r = 0.98, p < 0.001; Fig. 3) showed an effect in the opposite direction to playback (downregulated in heat-call embryos while upregulated in females), implying that the sex imbalance would tend to attenuate rather than inflate playback-associated differences. While we did not recover significant GO-term enrichment for the brown module, the strong module–sex association and the presence of sex-linked transcripts among sex-by-treatment interactions (e.g., Z-linked *HDHD2* and W-linked *LOC116806857* and *LOC116806879*/*KCMF1*) still support sex-biased regulation at this developmental stage. Consistent with this, future studies with balanced sex ratios are likely to improve power to detect additional playback-responsive transcripts and to more fully resolve sex-by-treatment interactions (*HDHD2*, *LOC116806857*, *LOC116806879*/*KCMF1*, *ABRACL*, *PTPRM*).

## 4 Conclusion

Here we showed that an acoustic signal of heat from the parents is capable of triggering changes to offspring BBB and mural cells that resemble the first response of the brain to heat exposure, and the prolongation of a plastic vascular phenotype during development (Bain et al., 2015; Boorman et al., 2023; Huber et al., 2001). The evidence suggest that these changes correspond to a localized brain defense mechanism against upcoming heat-challenges, signaled by heat-calls during incubation. Our study therefore hints at a so-far unforeseen mechanism of heat adaptation in avian embryos and extends the impact of prenatal sound on development. Likewise in a recent study, we showed that light exposure, that uses another sensory modality, causes gene expression changes in the vascular and circulatory pathways of embryonic chick retina (Versace et al., 2022). Together, these findings align with recent developments in medical research pointing at the high plasticity of neurovascular cells and the BBB, and their potential importance for developmental programming (Phan et al., 2025; Zhao et al., 2022). This is particularly important in the context of accelerating climate change (Intergovernmental Panel on Climate Change, 2015; Stillman, 2019; Ummenhofer & Meehl, 2017), with heat-related mortality projected to increase not only in wildlife, including arid-adapted species like the zebra finch (Albright, 2017; Conradie, 2020), but also in humans, where heatwaves pose escalating public health hazard (Howard et al., 2024; Narayanan & Keellings, 2025; World Health Organization, 2024). Our findings also open an intriguing possibility for future therapeutic application, using sound (heat-call) exposure as a safer alternative to the use of short heat exposure for transiently opening the blood-brain barrier for drug delivery to the brain. Overall, our study opens unique research opportunities on brain heat-adaptation, sensory-mediated developmental plasticity and beyond.

## 5 Methods

### 5.1 Experimental Design

To obtain eggs for playback treatment, adult zebra finches were allowed to pair and breed in an outdoor aviary at Deakin University, Australia. Eggs collected on the laying day were placed in fan-less artificial incubator (Bellsouth 100 electronic incubator) at 37.5°C for nine days and 60% humidity. On day 10 (E10) in the morning, the eggs were randomly allocated, using a within clutch design and balancing laying order, to one of two experimental incubators, each equipped with a different acoustic playback treatment: control call playback (whine calls combined with tet calls, n = 9) or heat call playback (whine calls combined with heat calls, n = 10). We included 9–10 biological replicates per treatment group, which is comparable to, or exceeds sample sizes commonly reported in adult zebra finch hypothalamus studies (n = 4–6 per group) (Kumari et al., 2022). Whine calls are contact calls with complex acoustic structure that may contribute to auditory system development, while tet calls are typical contact calls commonly uttered by parents at the nest but not related to temperature (Boucaud et al., 2016). By contrast, heat calls are fast, high-pitched vocalizations produced by adult zebra finches under thermal stress (Pessato et al., 2020). Prenatal acoustic playbacks were broadcast daily from E10 to E13 for 9 hours per day (10:00–19:00) at 65 dB sound pressure level. Playbacks were delivered via speakers (Sennheiser HD439, Video Guys, Melbourne, Australia) positioned inside each experimental incubator and externally connected to an amplifier (Digitech 18 W, Jaycar, Geelong, Australia) and audio player (Zoom H4nSP and Marantz PMD670, Video Guys, Melbourne, Australia). To control for potential incubator-specific effects, egg trays and audio playback equipment were swapped daily between the two experimental incubators. On the night of E13, embryos from both playback treatments were transferred to a single shared silent incubator to equalize the immediate pre-sampling environment.

On the morning of E14 (i.e., day before hatch) between 10:00 and 12:00, embryos were extracted from eggs and euthanized by rapid decapitation. Isolated heads were immediately embedded in optimal cutting temperature (OCT) compound and flash-frozen on dry ice. Frozen tissue blocks were shipped on dry ice from Deakin University (Australia) to the Queen Mary University of London (United Kingdom) and stored at −80°C upon arrival until cryosectioning for hypothalamic tissue collection. Brain tissue was sectioned coronally at 100 μm thickness on a cryostat, and a single cylindrical punch (1 mm diameter) was collected when the third ventricle became visible, approximately 5,000 μm caudal to the beak (Supplemental Methods). Prior to sampling experimental embryos, we validated that punches captured hypothalamic tissue by performing fluorescent in situ hybridization with hypothalamic gene markers in whole brain coronal sections of untreated embryos (Supplemental Methods and Fig. S3). Tissue punches were immediately collected into cryotubes on dry ice and stored at −80°C until RNA extraction. All procedures were approved by the Animal Ethics Committee of Deakin University (G29-2016) and conducted in accordance with Australian guidelines for animal research.

### 5.2 RNA Extraction, Sequencing, and Statistical Analysis

Total RNA was extracted using the Zymo Quick-RNA Microprep Kit (cat #R1050) at the Genome Centre, Queen Mary University of London. RNA quality and concentration were assessed using NanoDrop spectrophotometry and Agilent 2100 Bioanalyzer. The RNA-seq experiment was conducted on the Illumina NextSeq 500 platform in 75 base pair, paired-end mode. Quality control was performed by trimming low quality sequences using Trimmomatic (v 0.39) (Bolger et al., 2014). The RNA-seq reads were mapped to the zebra finch reference transcriptome (bTaeGut1.4.pri) and the transcripts were quantified using the R package Salmon (v 1.5.2) (Patro et al., 2017). A total of 49,298 transcripts corresponding to 21,407 genes were detected and quantified across all samples.

Gene-level counts were normalized and modeled using R package DESeq2 (v 1.30.1) (Love et al., 2014). Genes with less than 10 counts in total were excluded. All 19 embryos passed quality control and were included; no samples were excluded post hoc. The primary model tested the main effect of treatment (∼ sex + playback condition) using a likelihood ratio test (LRT) against a reduced model containing only sex. Log2 fold changes were shrunk using apeglm to stabilize variance estimates and improve ranking. Significant differentially expressed genes (DEGs) were defined as those with Benjamini–Hochberg adjusted p-value (p_adj_) < 0.05. The final set comprised 49 DEGs (48 downregulated, 1 upregulated). Secondary analysis assessed the interaction term (∼ sex + playback condition + sex:playback condition) to investigate sex-by-treatment interaction at p_adj_ < 0.05.

Principal component analysis (PCA) was performed on variance-stabilization transformed (VST) data to characterize total expression variance.

The differential expression analyses in DESeq2 used negative binomial generalized linear models with gene-wise dispersion estimates and mean-variance trends fitted from the data. All p-values obtained from the Wald and likelihood ratio tests were adjusted for multiple comparisons using the Benjamini–Hochberg false discovery rate (FDR) method. VST was applied to normalized counts to obtain approximately homoscedastic, log-like expression values for principal component analysis and downstream correlation-based analyses (see section 5.4).

To complement the genome-wide approach, we performed a targeted differential expression analysis on a literature-derived set of 143 genes with established hypothalamic function (Table S1). This gene list was compiled from published expression atlases and functional studies of the avian and mammalian hypothalamus (Antin et al., 2007; Bell et al., 2004; Lovell et al., 2020; Rouillard et al., 2016; Uhlén et al., 2015). The full DESeq2 dataset (19,137 genes) was subset to include only those hypothalamic genes present in the filtered count matrix. Differential expression was then tested on this reduced gene set using DESeq2 with the same model specification (∼ sex + playback condition) and likelihood ratio test framework as the genome-wide analysis. Log2 fold changes were estimated using apeglm shrinkage. Multiple testing correction was applied via the Benjamini–Hochberg method within the hypothalamic gene subset. This targeted approach reduces the multiple-testing burden relative to the whole-transcriptome analysis, thereby increasing sensitivity to detect modest expression changes in genes of a priori biological interest.

To identify biological pathways and functional categories associated with heat call exposure, DEGs were subjected to Gene Ontology (GO) and KEGG pathway enrichment analysis for biological processes using ShinyGO v0.85.1 (species: *Homo sapiens*) (Ge et al., 2020). To mitigate incomplete functional annotation in the zebra finch genome, we mapped zebra finch gene symbols to human orthologs using the biomaRt R package (v 2.60.1) to query the Ensembl BioMart database (Ensembl release 110, *Taeniopygia guttata* genome assembly GCF_003957565.2_bTaeGut1.4.pri) (Kinsella et al., 2011). Zebra finch genes with identifiable human orthologs were used as input for functional enrichment analysis. To ensure the statistical universe matched the set of genes measurable in our experiment, we provided a custom background gene set consisting of all 21,407 zebra finch genes included in the RNA-seq analysis that successfully mapped to human orthologs (N = 11,310 unique human genes). Enrichment was tested against GO Biological Process, GO Molecular Function, GO Cellular Component, and KEGG pathway databases. Enrichment P-values were calculated using the hypergeometric test and corrected for multiple testing using the Benjamini–Hochberg false discovery rate (FDR) method. Pathways meeting FDR < 0.05 were considered statistically significant. Pathway size limits were set to a minimum of 2 and a maximum of 5,000 genes. Redundancy reduction was enabled to collapse highly overlapping pathways (95% gene overlap and 50% name overlap). Results were ranked by fold enrichment within the FDR-significant set, and the top pathways per module were reported as bar plots displaying fold enrichment and FDR values.

### 5.3 Isoform Switch Analysis

To identify playback condition-wise splice-isoform usage patterns in our RNA-Sequencing data, we used IsoformSwitchAnalyzeR package in R (v2.4.0) to perform an isoform switching analysis (Vitting-Seerup & Sandelin, 2019). A design matrix with sound playback conditions and sex as covariates was constructed from sample metadata (n=19). We identified transcripts at the isoform resolution by using a gtf and transcript FASTA files (bTaeGut1.4.pri assembly). We filtered the transcript isoforms of genes by removing genes with (i) expression <1 Transcript per Million (TPM), (ii) isoforms with 0 TPM, and (iii) single-isoform genes. Differential isoform usage was quantified using DEXSeq (α=0.05 FDR-adjusted; differential isoform fraction ≥ 0.05) using the isoformSwitchTestDEXSeq function. We characterized splicing events using two complementary approaches: at the genome level (via extractSplicingGenomeWide) and at the event or mechanistic level (via extractSplicingEnrichment). We used the default junction overlap thresholds to calculate intron retention frequencies in transcripts using the analyzeAlternativeSplicing function. Finally, changes in isoform usage and expression between control and heat call playback conditions were visualized.

### 5.4 Weighted Gene Co-expression Network Analysis (WGCNA)

To identify patterns of gene co-expression and their relationships with different embryonic sound playbacks, we conducted a Weighted Gene Co-expression Network Analysis (WGCNA) using the WGCNA (v 1.73) package in R (Langfelder & Horvath, 2008). The input transcript read counts were prepared based on the zebra finch reference transcriptome (GCF_003957565.2_bTaeGut1.4.pri assembly) using Salmon version 1.5.2 (Patro et al., 2017). These gene expression raw counts were normalized using the variance stabilizing transformation (VST) method in the DESeq2 package. A DESeq2 dataset was constructed with intercept only in the design formula. The VST normalization was applied ignoring the experimental design to produce variance stabilized, log-like expression values. The resulting normalized expression matrix was transposed in a sample-by-genes format for downstream analyses. Networking construction parameters like the soft-thresholding power were selected using an iterative approach. The scale-free topology fit index (R^2^) and mean connectivity were calculated for a range of powers from 1 to 30 to find the optimal power (power = 8) achieving a high scale-free topology fit. The network was constructed using a minimum module size of 30 genes and a module similarity (i.e., merge cut height) of 0.25.

To examine module–trait relationships, module assignments were converted to color labels, module eigengenes were calculated, and Pearson correlations were computed between eigengenes and sex or playback condition using VST-normalized expression values. Significance of these correlations was assessed using two-sided Student’s t-tests, and results were visualized as a heatmap annotated with correlation coefficients and corresponding p-values. We performed a Gene Ontology (GO) and KEGG pathway enrichment analysis as described previously (see section 5.2) for modules significantly correlated with heat call playback. For each module, genes with identifiable human orthologs were used as input for functional enrichment analysis. Results were reported as bar plots displaying fold enrichment and FDR values.

To find the hub genes in each module, we calculated the intramodular connectivity (kWithin) for each gene using the adjacency-matrix raised to the selected-threshold power. For the green module, edges with adjacency weights below 0.4 were excluded to focus on robust co-expression patterns. The top five hub genes, with the highest kWithin calculated from the adjacency matrix, were identified for the green module. These hub genes and their associated neighbors were visualized as network layouts using the Kamada-Kawai algorithm using the ggraph package (v 2.2.1) (Pedersen, 2024).

### 5.5 Cell-Type Enrichment Analysis

To establish cellular specificity of the identified modules from the WGCNA analysis, we conducted a cell-type enrichment analysis using a publicly available dataset “HYPOMAP: A comprehensive spatio-cellularmap of the human hypothalamus” available at the Chan Zuckerberg CELLxGENE Discover database (Megill et al., 2021; Tadross et al., 2025). A MAST test was conducted using the FindAllMarkers function in the Seurat R package (v 5.2.1) (Hao et al., 2024) to identify the top 100 cell-type marker genes with minimum expression in 25% of cells and a minimum log fold-change threshold of 0.25. These marker genes were converted from ensembl identifiers to human gene symbols using the org.Hs.eg.db annotations (Carlson, 2024). Next, the zebra finch genes co-expressed in the WGCNA green module were mapped to their human orthologs using the biomaRt R package (v 2.60.1) interfaced with Ensembl (release 110, accessed July 2025) (Kinsella et al., 2011). Finally, cell-type-module associations were quantified using a hypergeometric test by comparing each module’s gene set against cell type markers. The p-values were adjusted for multiple-testing correction using the Benjamini-Hochberg method and the results were visualized using a heatmap.

### 5.6 Deconvolution Analysis

To estimate the cellular composition of our bulk-tissue RNA-Seq data and obtain cell-type-specific abundance, we used CIBERSORTx (v 1.0) (Newman et al., 2019). Bulk tissue mixture files containing transcript-level normalized counts (counts per million or CPM) were prepared according to the package’s recommendations. A custom signature matrix was generated from a single-cell RNA-sequencing atlas of the human hypothalamus (HYPOMAP: A comprehensive spatio-cellular map of the human hypothalamus) available at the Chan Zuckerberg CELLxGENE Discover database (Megill et al., 2021; Tadross et al., 2025). The cell fractions mode was used to construct a batch-corrected signature matrix and adjusted bulk mixture files with 500 permutations and without quantile normalization using the Docker version of CIBERSORTx. These batch-corrected files were subsequently uploaded to the CIBERSORTx web portal to run a custom cell fractions analysis with 1000 permutations in absolute mode, and the output files were visualized using R.

CIBERSORTx was run in absolute mode, which reports RNA-based abundance scores for each cell type that scale with an overall "absolute score" per sample and are therefore not constrained to sum to 1. To derive within-sample cell-type proportions, absolute-mode scores were normalized by dividing each cell-type value by the sample’s absolute score, such that relative fractions sum to 1 across the cell types represented in the signature matrix. For each sample, CIBERSORTx reported a goodness-of-fit p-value, the Pearson correlation between observed and model-predicted bulk expression, and the root mean square error (RMSE). In this dataset, all samples had p < 0.0001, Pearson r > 0.89, and RMSE between 0.70 and 0.75 (Table S6).

### 5.7 Use of AI Tools

Large language models accessed via the Perplexity platform (including Grok 4.1, GPT 5.1, Sonar, and Claude Sonnet 4.5) were employed to assist with R script optimization and debugging of computational workflows, including RNA-seq, WGCNA, isoform switching, and deconvolution pipelines. These models were also used to help edit and refine portions of the manuscript, particularly the Materials and Methods and figure legends, to ensure adherence to the target journal’s formatting and reporting guidelines. All AI-assisted code and text were critically reviewed, modified, and validated prior to inclusion in the manuscript, and the authors take full responsibility for the accuracy and integrity of all content. No AI tools were used for image creation, editing, or enhancement.

## 7 Author Contributions

P.S. conceived the bioinformatic analyses, performed all computational and statistical analyses, generated figures and tables, and led the drafting and revision of the manuscript. K.P. performed embryonic dissections, hypothalamic punching, and tissue collection for RNA-seq, and contributed to manuscript editing. M.G.E. performed embryonic dissections, hypothalamic punching, and tissue collection for RNA-seq, and contributed to data interpretation and manuscript editing. E.V. contributed to conceptualization and experimental design, supervised aspects of laboratory work, managed funding, and contributed to data interpretation and manuscript editing. K.L.B. contributed to conceptualization and experimental design, supervised animal husbandry and playback experiments, contributed to data interpretation, obtained funding, and co-wrote and edited the manuscript. M.M.M. contributed to conceptualization and experimental design, animal work and playback exposure, contributed to data interpretation, obtained funding, and co-wrote and edited the manuscript. D.F.C. contributed to conceptualization, experimental design, and interpretation of gene expression analyses, secured funding, and co-wrote and edited the manuscript. J.M.G. conceived and supervised the overall project, contributed to experimental design, data analysis, and interpretation, obtained funding, and co-wrote and edited the manuscript. All authors read and approved the final manuscript.

## Acknowledgments

The authors thank the animal care staff and technical personnel at Deakin University for assistance with zebra finch husbandry and incubation logistics. RNA sequencing was supported by the Genome Centre and Queen Mary University of London, and we thank its staff for expert technical support.

We are grateful to the Clemson Light Imaging Facility (CLIF) for access to imaging resources and for substantial assistance with fluorescence microscopy and image acquisition. Rhonda Powell as well as members, management, and staff at CLIF provided valuable technical guidance and intellectual input related to imaging of embryonic brain tissue.

## **9** Ethics Approval

All animal procedures were approved by the Animal Ethics Committee of Deakin University (G29-2016) and conducted in accordance with Australian guidelines.

## **10** Data Accessibility and Benefit-Sharing Section

### 10.1 Data Accessibility Statement

Raw RNA-seq data and processed gene expression files have been deposited in NCBI’s Gene Expression Omnibus (GEO) and are publicly accessible through GEO Series accession number GSE313742 (https://www.ncbi.nlm.nih.gov/geo/query/acc.cgi?acc=GSE313742). Raw sequencing reads are also available through the NCBI Sequence Read Archive (SRA) under BioProject accession PRJNA1381516 (https://www.ncbi.nlm.nih.gov/bioproject/PRJNA1381516). Sample metadata and count matrices are available as supplementary files with the GEO submission. Intermediate analysis files and all computational analysis code, including R scripts for differential expression, WGCNA, isoform switching, and deconvolution analyses, are publicly available on GitHub (https://github.com/praxsubba/Heat_Call_Reprogramming_Bulk_RNA_Seq). All other relevant data, including supplementary tables and figures, are provided within the article and its supplementary information.

### **10.2** Benefit-Sharing Statement

This research was conducted under an international scientific collaboration among researchers from Australia, the United Kingdom, Spain, and the United States. All collaborating scientists are included as co-authors and contributed meaningfully to the research design, execution, and interpretation as detailed in the Author Contributions section. Research findings have been shared with all collaborating institutions and will be disseminated to the broader scientific community through open-access publication and publicly archived datasets (see Data Accessibility Statement above). All biological samples were collected under appropriate ethical approvals from Australian authorities (Deakin University Animal Ethics Committee (G29-2016), and the research complies with the Convention on Biological Diversity and relevant national laws. Zebra finches used in this study were captive-bred populations maintained at Deakin University, and no wild populations or endangered species were sampled. Benefits from this research accrue from the open sharing of data, analytical code, and results as described above, contributing to the understanding of developmental programming mechanisms relevant to climate change adaptation.

