## Supplemental Data for "Programming of Embryonic Blood Brain Barrier and Neurovascular Transcriptome by an Anticipatory Acoustic Signal of Heat in the Zebra Finch"


### Abstract

Organisms have evolved mechanisms to adjust to rapid environmental change. A dramatic example is the Australian zebra finch, where incubating parents produce an acoustic signal ("heat call") during extreme heat, triggering adaptive phenotypic plasticity in their offspring growth, thermoregulation, and reproductive success. To elucidate for the first time the molecular mechanisms underlying heat call-induced programming, we hypothesized a prenatal shift in hypothalamic gene expression, given the hypothalamus's central role in neuroendocrine signaling controlling metabolism and growth. In addition, we tested whether prenatal heat-call exposure induces local changes in the brain, to protect this highly heat-sensitive organ from upcoming heat challenges. We exposed zebra finch embryos to chronic playback of parental heat calls or control calls, then isolated the hypothalamus for RNA sequencing to identify differentially expressed genes and gene regulatory networks. Heat-call exposure elicited modest neuroendocrine gene expression changes, but robust downregulation of genes tied to muscle contraction and cytoskeletal dynamics, with evidence of isoform usage shifts. These changes were prominently localized to hypothalamic neurovascular endothelial, mural, and ependymal cell populations, forming the blood–brain barrier (BBB). Because embryos experienced heat-associated sound, but not heat itself, and changes matched the loosening of the BBB to avoid breakages, these transcriptomic shifts likely represent an anticipatory response to enhance subsequent brain resilience to heat. Our study provides the first genome-wide characterization of embryonic hypothalamic gene expression in a songbird and reveals that prenatal acoustic cues can developmentally program neurovascular systems, expanding current understanding of developmental plasticity under climate change.


### Supplementary Methods

#### Methods for Hybridization Chain Reaction

To validate the presence of hypothalamic tissue in the regions surrounding the third ventricle, we performed hybridization chain reaction (HCR) for the *SIM1* gene (single-minded bHLH transcription factor 1), a marker for the paraventricular nucleus (PVN) identified from single-nucleus RNA sequencing data of zebra finch hypothalamus (unpublished results). Coronal sections of E13 embryos from three untreated animals were collected caudal to the eye, where the third ventricle becomes visible. Brains were sectioned at 20 µm using a Leica CM3050 S cryostat (Leica Microsystems, Wetzlar, Germany) and mounted on Superfrost Plus slides. Tissue sections were immediately fixed in 4% paraformaldehyde, hydrated in 1× PBS, dehydrated through a 70%, 95% and 100% ethanol series, and stored at −20°C until the following day for hybridization.

Fixed tissue sections were processed for HCR™ Gold RNA-FISH following the manufacturer's sample-on-slide protocol, with minor modifications (Choi et al., 2010, 2014, 2016, 2018). Notably, samples were thawed and rehydrated before probe hybridization; the protocol was optimized by extending probe incubation to overnight and amplification to 3 h. Sections were then co-stained with DAPI (1 µg $ml^{-1}$) and imaged using the Leica SR GSD widefield super-resolution microscope with TIRF module (Leica Microsystems, Wetzlar, Germany).

Fluorescence images were acquired as single-plane TIFF files using Leica LAS X software on the Leica SR GSD widefield microscope (no Z-stacks or deconvolution were applied). Raw TIFF images were imported into FIJI/ImageJ (v2.16.0/1.54p) and converted to 16-bit grayscale prior to visualization (Schindelin et al., 2012). For display, individual channels (*SIM1* probe signal and DAPI) were pseudocoloured (*SIM1*: magenta; DAPI: cyan) and merged to generate composite images. Only linear adjustments to brightness and contrast were applied uniformly across each image, and no further image processing, filtering, or quantitative analysis was performed.

### Supplementary Figures and Tables

### Supplementary Figures

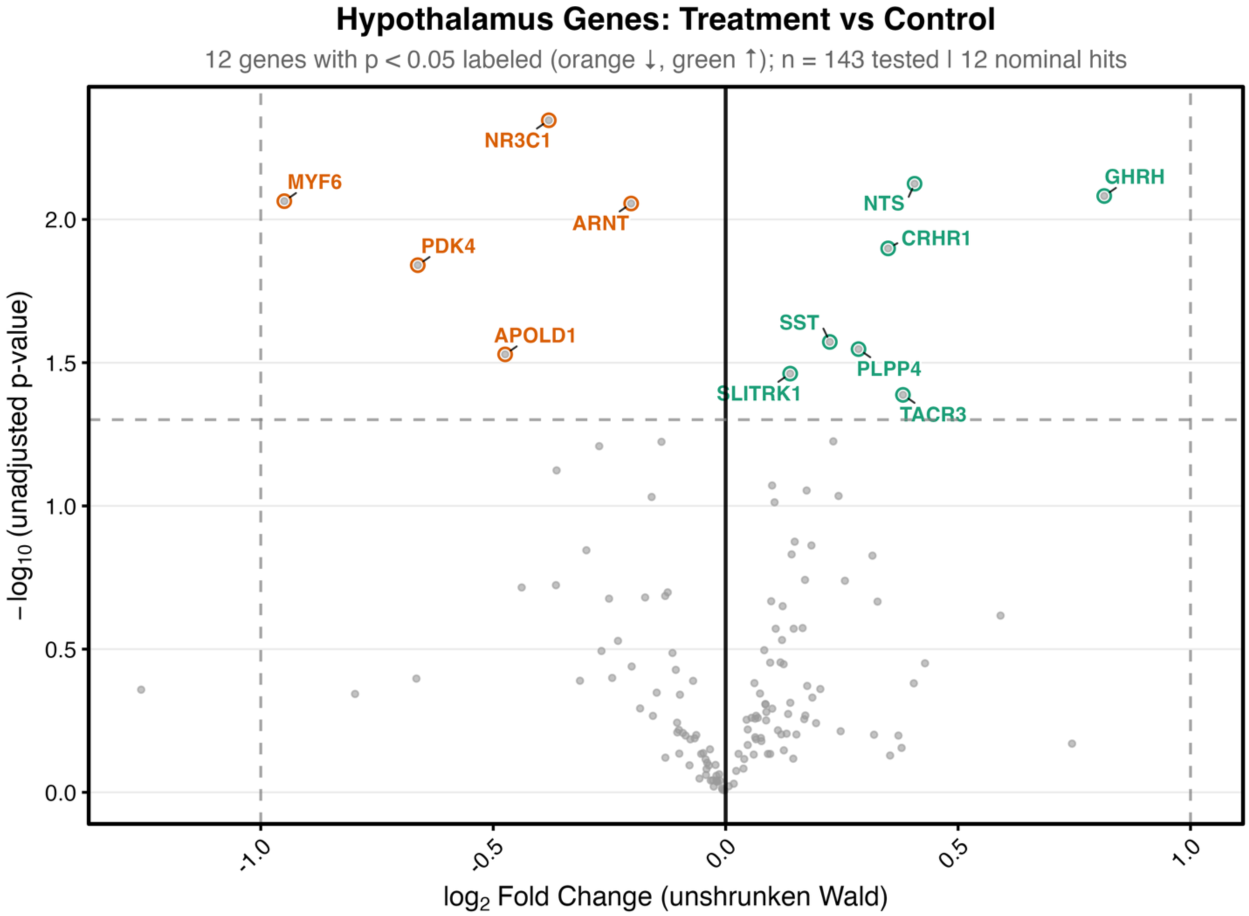


**Fig. S1: Results of the Targeted Differential Gene Expression Analysis:** Targeted differential gene expression analysis of 143 hypothalamic genes in medial hypothalamic punches (n=19: 10 heat call, 9 control embryos). Volcano plot shows log2 fold change (unshrunken Wald estimates; x-axis) versus −log10(unadjusted p-value; y-axis) from DESeq2. Twelve genes with nominal significance ($p < 0.05$, padj > 0.25) are highlighted: orange (downregulated with negative apeglm-shrunken log2FC: *NR3C1, MYF6, ARNT, PDK4, APOLD1*) and green (upregulated: *NTS, GHRH, CRHR1, SST, PLPP4, SLITRK1, TACR3*). Dashed lines: log2FC = ±1, p = 0.05.

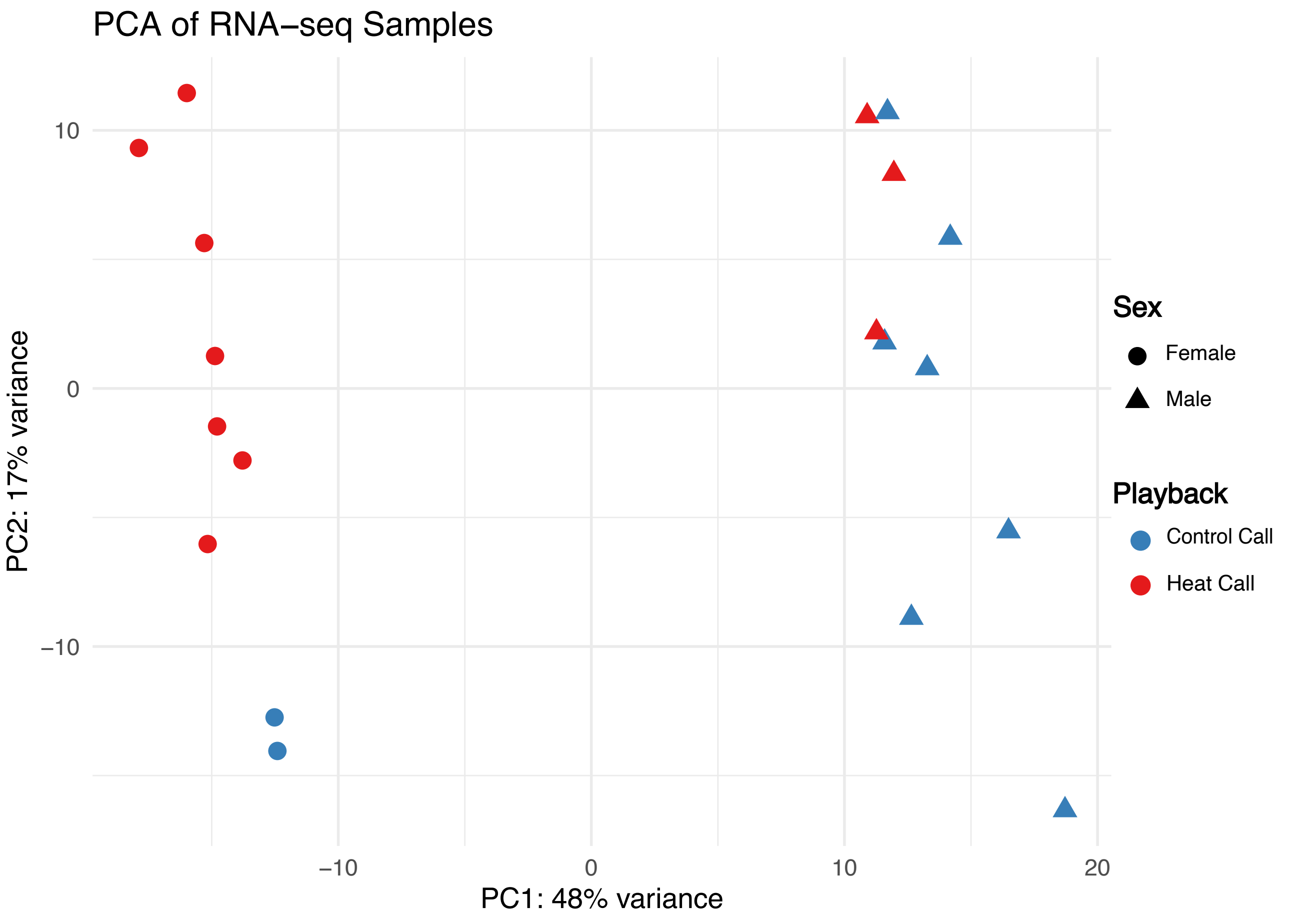


**Fig. S2:** Principal component analysis of variance-stabilizing transformed (VST) count data from medial hypothalamic punches (n=19 biological replicates: 10 heat call, 9 control). PC1 (48% variance) separates samples by sex; PC2 (17% variance) captures additional variation, including playback-associated separation Circles = female, triangles = male; red = heat call, blue = control call.

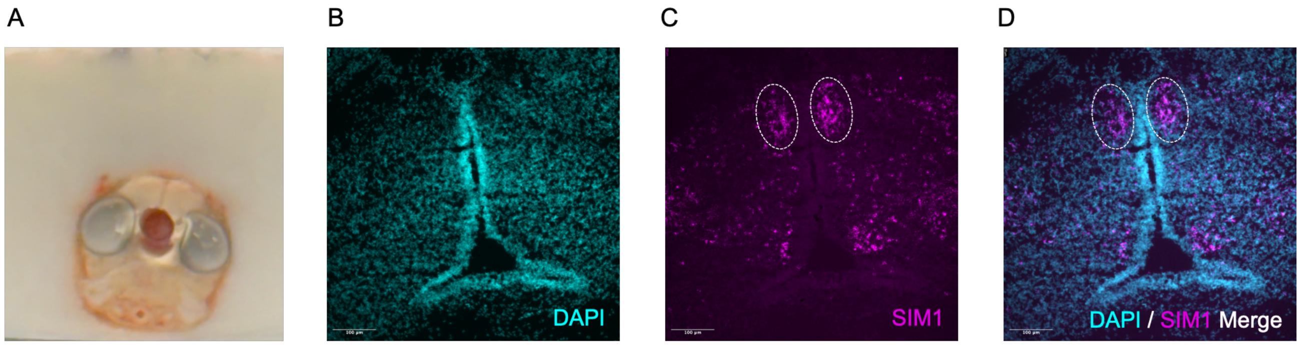


**Fig. S3:** (A) Medial hypothalamic punch encompassing the third ventricle and various hypothalamic nuclei from E13 Embryo. (B-D) Fluorescence in situ hybridization (FISH) pilot validation experiment of SIM BHLH Transcription Factor 1 (*SIM1*) expression in the developing hypothalamus. Representative image showing the hypothalamic marker gene *SIM1* mRNA signal in the (C) paraventricular nucleus of the hypothalamus (magenta), located lateral to the third ventricle, in an untreated embryo. (B and D) Nuclei are counterstained with DAPI (cyan). (C and D) Dotted lines delineate *SIM1*-positive cells within the paraventricular nucleus.

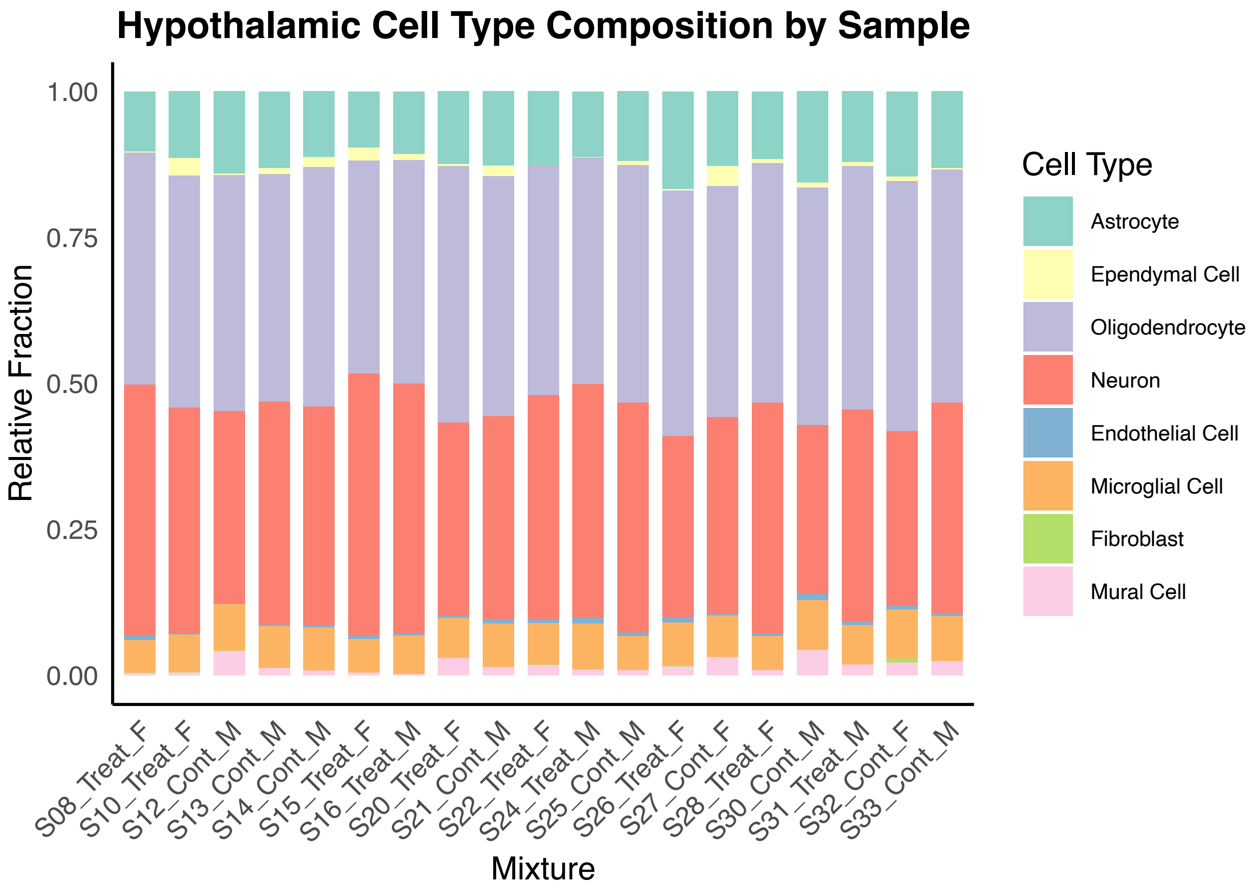


**Fig. S4:** CIBERSORTx deconvolution of bulk hypothalamic RNA-seq samples (n = 19: 10 heat call, 9 control; one medial punch per embryo). Stacked bars show estimated relative cell-type fractions (normalized to sum to 1) for eight brain cell types defined by the HYPOMAP human hypothalamus reference signature matrix. For each sample, CIBERSORTx reported a Monte Carlo permutation p-value (all samples $p < 0.0001$), the Pearson correlation coefficient between observed and model-predicted bulk expression restricted to signature genes ($r > 0.89$), and the root mean square error (RMSE; 0.70–0.75) between observed and model-predicted expression.

**Table S1: Hypothalamic Gene List for Targeted Analysis:** Targeted differential gene expression results for 143 literature-derived hypothalamic genes in medial hypothalamic punches from heat call and control embryos.

| Gene | Base Mean | Log2FC Unshrunken | LFC SE Unshrunken | Unadjusted p-value | FDR | Log2FC Shrunken apeglm | LFC SE Shrunken apeglm |
|---|---|---|---|---|---|---|---|
| ***NR3C1*** | 1640.00 | -0.38 | 0.13 | **0.00** | 0.25 | -0.26 | 0.17 |
| ***NTS*** | 1204.06 | 0.41 | 0.15 | **0.01** | 0.25 | 0.05 | 0.10 |
| ***GHRH*** | 72.15 | 0.81 | 0.31 | **0.01** | 0.25 | 0.02 | 0.05 |
| ***MYF6*** | 88.39 | -0.95 | 0.36 | **0.01** | 0.25 | -0.02 | 0.05 |
| ***ARNT*** | 1125.44 | -0.20 | 0.08 | **0.01** | 0.25 | -0.13 | 0.09 |
| ***CRHR1*** | 744.52 | 0.35 | 0.14 | **0.01** | 0.29 | 0.07 | 0.15 |
| ***PDK4*** | 193.41 | -0.66 | 0.27 | **0.01** | 0.29 | -0.02 | 0.05 |
| ***SST*** | 5416.36 | 0.22 | 0.10 | **0.03** | 0.42 | 0.07 | 0.11 |
| ***PLPP4*** | 5652.55 | 0.29 | 0.13 | **0.03** | 0.42 | 0.05 | 0.09 |
| ***APOLD1*** | 164.51 | -0.47 | 0.22 | **0.03** | 0.42 | -0.02 | 0.05 |
| ***SLITRK1*** | 8001.38 | 0.14 | 0.07 | **0.03** | 0.45 | 0.07 | 0.07 |
| ***TACR3*** | 177.28 | 0.38 | 0.19 | **0.04** | 0.49 | 0.02 | 0.05 |
| ***KCNA1*** | 871.28 | 0.23 | 0.12 | 0.06 | 0.59 | 0.03 | 0.06 |
| ***BMP7*** | 1452.55 | -0.14 | 0.07 | 0.06 | 0.59 | -0.06 | 0.07 |
| ***FOSL2*** | 672.56 | -0.27 | 0.15 | 0.06 | 0.59 | -0.03 | 0.06 |
| ***EMP1*** | 666.97 | -0.36 | 0.20 | 0.08 | 0.66 | -0.02 | 0.05 |
| ***ARPC5*** | 19648.00 | 0.10 | 0.06 | 0.08 | 0.66 | 0.05 | 0.05 |
| ***SLC10A4*** | 1052.07 | 0.17 | 0.10 | 0.09 | 0.66 | 0.04 | 0.06 |
| ***SLC18A3*** | 1403.33 | 0.24 | 0.14 | 0.09 | 0.66 | 0.02 | 0.05 |
| ***SLC2A1*** | 2260.96 | -0.16 | 0.09 | 0.09 | 0.66 | -0.04 | 0.06 |
| ***GCH1*** | 1813.70 | 0.11 | 0.06 | 0.10 | 0.66 | 0.05 | 0.05 |
| ***KCNJ6*** | 690.93 | 0.15 | 0.10 | 0.13 | 0.82 | 0.03 | 0.05 |
| ***PENK*** | 8758.52 | 0.18 | 0.12 | 0.14 | 0.82 | 0.03 | 0.05 |
| ***PTGS2*** | 168.03 | -0.30 | 0.20 | 0.14 | 0.82 | -0.02 | 0.05 |
| ***SIX3*** | 2676.85 | 0.14 | 0.10 | 0.15 | 0.82 | 0.03 | 0.05 |
| ***TRH*** | 195.78 | 0.32 | 0.22 | 0.15 | 0.82 | 0.01 | 0.05 |

| | | | | | | | |
|---|---|---|---|---|---|---|---|
| ***TAC1*** | 3141.34 | 0.17 | 0.13 | 0.18 | 0.86 | 0.02 | 0.05 |
| ***PVALB*** | 1263.46 | 0.26 | 0.19 | 0.18 | 0.86 | 0.01 | 0.05 |
| ***BMP4*** | 219.36 | -0.37 | 0.28 | 0.19 | 0.86 | -0.01 | 0.05 |
| ***MYH11*** | 180.52 | -0.44 | 0.34 | 0.19 | 0.86 | -0.01 | 0.05 |
| ***NOTCH1*** | 2013.09 | -0.12 | 0.10 | 0.20 | 0.86 | -0.02 | 0.05 |
| ***SGK1*** | 2121.87 | -0.13 | 0.10 | 0.21 | 0.86 | -0.02 | 0.05 |
| ***EDN3*** | 386.01 | -0.17 | 0.14 | 0.21 | 0.86 | -0.02 | 0.05 |
| ***THRA*** | 778.78 | -0.25 | 0.20 | 0.21 | 0.86 | -0.01 | 0.05 |
| ***SGSM1*** | 1288.55 | 0.10 | 0.08 | 0.22 | 0.86 | 0.03 | 0.05 |
| ***GAL*** | 764.59 | 0.33 | 0.26 | 0.22 | 0.86 | 0.01 | 0.05 |
| ***RASGRF2*** | 757.89 | 0.12 | 0.10 | 0.22 | 0.87 | 0.02 | 0.05 |
| ***LOC100219863*** | 106.26 | 0.59 | 0.50 | 0.24 | 0.91 | 0.01 | 0.05 |
| ***CBLN2*** | 4674.73 | 0.17 | 0.15 | 0.27 | 0.91 | 0.01 | 0.05 |
| ***NGB*** | 345.51 | 0.15 | 0.13 | 0.27 | 0.91 | 0.01 | 0.05 |
| ***ODC1*** | 2999.70 | 0.11 | 0.10 | 0.27 | 0.91 | 0.02 | 0.05 |
| ***TAGLN3*** | 1765.27 | 0.12 | 0.12 | 0.29 | 0.91 | 0.02 | 0.05 |
| ***DRD2*** | 179.96 | -0.23 | 0.22 | 0.30 | 0.91 | -0.01 | 0.05 |
| ***ATP2B2*** | 7999.47 | 0.08 | 0.08 | 0.32 | 0.91 | 0.03 | 0.05 |
| ***S100A10*** | 1583.99 | -0.27 | 0.27 | 0.32 | 0.91 | -0.01 | 0.05 |
| ***PTCH1*** | 1941.38 | -0.11 | 0.12 | 0.33 | 0.91 | -0.02 | 0.05 |
| ***CALB2*** | 5086.41 | 0.12 | 0.13 | 0.35 | 0.91 | 0.01 | 0.05 |
| ***ASCL1*** | 421.67 | 0.10 | 0.10 | 0.35 | 0.91 | 0.02 | 0.05 |
| ***NPVF*** | 434.31 | 0.43 | 0.46 | 0.35 | 0.91 | 0.00 | 0.05 |
| ***FGF10*** | 216.93 | 0.12 | 0.14 | 0.36 | 0.91 | 0.01 | 0.05 |
| ***SIX6*** | 610.80 | -0.20 | 0.22 | 0.36 | 0.91 | -0.01 | 0.05 |
| ***NUCB2*** | 5085.61 | -0.11 | 0.12 | 0.37 | 0.91 | -0.01 | 0.04 |
| ***HES5*** | 192.74 | -0.24 | 0.29 | 0.40 | 0.91 | -0.01 | 0.05 |
| ***NEUROD4*** | 10.16 | -0.67 | 0.79 | 0.40 | 0.91 | -0.00 | 0.05 |
| ***SLC38A8*** | 87.44 | -0.31 | 0.38 | 0.41 | 0.91 | -0.00 | 0.05 |
| ***IGF1R*** | 4671.41 | -0.07 | 0.08 | 0.41 | 0.91 | -0.02 | 0.04 |
| ***SDC3*** | 18110.13 | 0.06 | 0.08 | 0.42 | 0.91 | 0.02 | 0.04 |
| ***AVPI1*** | 66.14 | 0.40 | 0.50 | 0.42 | 0.91 | 0.00 | 0.05 |

| | | | | | | | |
|---|---|---|---|---|---|---|---|
| ***LOC100231196*** | 1935.95 | 0.18 | 0.22 | 0.43 | 0.91 | 0.01 | 0.05 |
| ***HDC*** | 103.68 | 0.20 | 0.26 | 0.44 | 0.91 | 0.01 | 0.05 |
| ***UTS2*** | 1.08 | -1.26 | 1.62 | 0.44 | 0.91 | -0.00 | 0.05 |
| ***BMP3*** | 1637.49 | -0.15 | 0.20 | 0.45 | 0.91 | -0.01 | 0.05 |
| ***CELF6*** | 3477.58 | 0.07 | 0.10 | 0.45 | 0.91 | 0.01 | 0.04 |
| ***LOC115491955*** | 1.44 | -0.80 | 1.06 | 0.45 | 0.91 | -0.00 | 0.05 |
| ***BMP2*** | 154.66 | -0.10 | 0.13 | 0.46 | 0.91 | -0.01 | 0.05 |
| ***TCF7L2*** | 4662.80 | 0.19 | 0.26 | 0.47 | 0.91 | 0.00 | 0.05 |
| ***CRH*** | 321.35 | 0.14 | 0.20 | 0.49 | 0.91 | 0.01 | 0.05 |
| ***TRHDE*** | 2164.43 | 0.09 | 0.12 | 0.49 | 0.91 | 0.01 | 0.04 |
| ***SCG3*** | 15178.33 | 0.09 | 0.13 | 0.49 | 0.91 | 0.01 | 0.04 |
| ***AOAH*** | 47.35 | -0.18 | 0.28 | 0.51 | 0.91 | -0.01 | 0.05 |
| ***SNCG*** | 2783.04 | 0.10 | 0.15 | 0.51 | 0.91 | 0.01 | 0.05 |
| ***SLC18A2*** | 1235.41 | 0.09 | 0.14 | 0.52 | 0.91 | 0.01 | 0.04 |
| ***NKX2-1*** | 812.18 | 0.13 | 0.22 | 0.53 | 0.91 | 0.01 | 0.05 |
| ***TRHR*** | 209.91 | 0.17 | 0.28 | 0.54 | 0.91 | 0.00 | 0.05 |
| ***FABP7*** | 95641.52 | 0.07 | 0.11 | 0.54 | 0.91 | 0.01 | 0.04 |
| ***CHRDL1*** | 717.28 | -0.16 | 0.26 | 0.54 | 0.91 | -0.01 | 0.05 |
| ***SSTR1*** | 7232.16 | 0.06 | 0.09 | 0.55 | 0.91 | 0.01 | 0.04 |
| ***RXFP3*** | 533.87 | 0.07 | 0.12 | 0.55 | 0.91 | 0.01 | 0.04 |
| ***PGRMC2*** | 2536.35 | 0.06 | 0.11 | 0.55 | 0.91 | 0.01 | 0.04 |
| ***PRSS56*** | 31.00 | 0.17 | 0.29 | 0.56 | 0.91 | 0.00 | 0.05 |
| ***LZTS3*** | 2261.80 | 0.04 | 0.08 | 0.56 | 0.91 | 0.01 | 0.04 |
| ***CRHR2*** | 1030.89 | 0.09 | 0.15 | 0.56 | 0.91 | 0.01 | 0.04 |
| ***ATP2A3*** | 711.20 | -0.10 | 0.18 | 0.57 | 0.91 | -0.01 | 0.05 |
| ***POU3F1*** | 149.97 | 0.19 | 0.35 | 0.57 | 0.91 | 0.00 | 0.05 |
| ***IRS4*** | 1706.76 | 0.05 | 0.09 | 0.60 | 0.91 | 0.01 | 0.04 |
| ***IGF1*** | 437.02 | -0.10 | 0.19 | 0.61 | 0.91 | -0.01 | 0.05 |
| ***RNF157*** | 4351.07 | 0.11 | 0.22 | 0.61 | 0.91 | 0.00 | 0.05 |
| ***MOV10L1*** | 58.69 | 0.25 | 0.49 | 0.61 | 0.91 | 0.00 | 0.05 |
| ***CALCB*** | 886.12 | -0.10 | 0.21 | 0.62 | 0.91 | -0.01 | 0.05 |
| ***TBX2*** | 481.88 | -0.09 | 0.19 | 0.62 | 0.91 | -0.01 | 0.05 |

| | | | | | | | |
|---|---|---|---|---|---|---|---|
| ***CARTPT*** | 761.33 | 0.13 | 0.27 | 0.62 | 0.91 | 0.00 | 0.05 |
| ***SYF2*** | 3449.48 | 0.12 | 0.25 | 0.63 | 0.91 | 0.00 | 0.05 |
| ***TSHR*** | 163.93 | 0.15 | 0.31 | 0.63 | 0.91 | 0.00 | 0.05 |
| ***LOC121469849*** | 87.79 | 0.32 | 0.66 | 0.63 | 0.91 | 0.00 | 0.05 |
| ***SSTR5*** | 339.33 | -0.06 | 0.13 | 0.63 | 0.91 | -0.01 | 0.04 |
| ***AVPR1B*** | 12.75 | 0.37 | 0.78 | 0.63 | 0.91 | 0.00 | 0.05 |
| ***MEST*** | 227.01 | -0.09 | 0.18 | 0.63 | 0.91 | -0.01 | 0.05 |
| ***INSIG1*** | 3892.42 | 0.06 | 0.14 | 0.64 | 0.91 | 0.01 | 0.04 |
| ***ROBO2*** | 5056.93 | 0.08 | 0.16 | 0.65 | 0.91 | 0.01 | 0.04 |
| ***NR3C2*** | 640.71 | -0.07 | 0.15 | 0.65 | 0.91 | -0.01 | 0.04 |
| ***MEIS1*** | 3783.81 | 0.07 | 0.14 | 0.65 | 0.91 | 0.01 | 0.04 |
| ***DLK1*** | 1271.10 | -0.08 | 0.17 | 0.65 | 0.91 | -0.00 | 0.04 |
| ***HMOX1*** | 1528.92 | 0.08 | 0.18 | 0.66 | 0.91 | 0.01 | 0.04 |
| ***PPP1R17*** | 2.00 | 0.75 | 1.78 | 0.68 | 0.91 | 0.00 | 0.05 |
| ***SHH*** | 1995.73 | 0.05 | 0.12 | 0.68 | 0.91 | 0.01 | 0.04 |
| ***CORT*** | 3.10 | 0.38 | 0.98 | 0.70 | 0.91 | 0.00 | 0.05 |
| ***DLL1*** | 868.77 | -0.03 | 0.09 | 0.71 | 0.91 | -0.01 | 0.04 |
| ***CHGA*** | 19735.64 | 0.13 | 0.34 | 0.71 | 0.91 | 0.00 | 0.05 |
| ***SCGN*** | 415.02 | -0.05 | 0.14 | 0.73 | 0.91 | -0.01 | 0.04 |
| ***OXTR*** | 153.68 | -0.10 | 0.29 | 0.73 | 0.91 | -0.00 | 0.05 |
| ***ERP29*** | 11960.98 | 0.03 | 0.08 | 0.73 | 0.91 | 0.01 | 0.04 |
| ***PDYN*** | 501.78 | 0.10 | 0.28 | 0.73 | 0.91 | 0.00 | 0.05 |
| ***ARHGAP35*** | 2783.63 | -0.05 | 0.15 | 0.73 | 0.91 | -0.00 | 0.04 |
| ***SSTR4*** | 575.92 | 0.09 | 0.27 | 0.73 | 0.91 | 0.00 | 0.05 |
| ***LHX5*** | 1669.55 | 0.06 | 0.18 | 0.74 | 0.91 | -0.00 | 0.04 |
| ***NHLH1*** | 4.63 | 0.35 | 1.08 | 0.74 | 0.91 | 0.00 | 0.05 |
| ***POMC*** | 1098.31 | -0.13 | 0.42 | 0.76 | 0.91 | -0.00 | 0.05 |
| ***SIM2*** | 44.89 | 0.15 | 0.48 | 0.76 | 0.91 | 0.00 | 0.05 |
| ***KCNIP2*** | 2067.67 | 0.04 | 0.13 | 0.77 | 0.91 | 0.00 | 0.04 |
| ***TH*** | 540.42 | -0.04 | 0.15 | 0.77 | 0.91 | -0.00 | 0.04 |
| ***AVPR1A*** | 365.60 | -0.04 | 0.15 | 0.79 | 0.93 | -0.00 | 0.04 |
| ***TRIP4*** | 893.79 | -0.02 | 0.09 | 0.80 | 0.93 | -0.00 | 0.04 |

| | | | | | | | |
|---|---|---|---|---|---|---|---|
| ***HMX2*** | 153.46 | -0.08 | 0.31 | 0.80 | 0.93 | -0.00 | 0.05 |
| ***SLC6A6*** | 5000.41 | -0.04 | 0.15 | 0.81 | 0.93 | -0.00 | 0.04 |
| ***SLC29A4*** | 1555.32 | 0.04 | 0.17 | 0.83 | 0.94 | 0.00 | 0.04 |
| ***SSTR2*** | 1650.08 | -0.04 | 0.19 | 0.83 | 0.94 | -0.00 | 0.04 |
| ***DBI*** | 9375.89 | 0.02 | 0.11 | 0.84 | 0.95 | 0.00 | 0.04 |
| ***HEY1*** | 1259.53 | -0.01 | 0.08 | 0.86 | 0.96 | -0.00 | 0.04 |
| ***SOX14*** | 1572.19 | -0.04 | 0.26 | 0.87 | 0.96 | -0.00 | 0.05 |
| ***THRB*** | 625.49 | -0.02 | 0.13 | 0.87 | 0.96 | -0.00 | 0.04 |
| ***LOC121469847*** | 824.27 | -0.06 | 0.43 | 0.89 | 0.96 | -0.00 | 0.05 |
| ***KCNMB4*** | 130.27 | -0.02 | 0.17 | 0.91 | 0.96 | -0.00 | 0.04 |
| ***FEZF1*** | 958.21 | -0.03 | 0.24 | 0.91 | 0.96 | -0.00 | 0.05 |
| ***SIM1*** | 124.53 | -0.03 | 0.28 | 0.91 | 0.96 | -0.00 | 0.05 |
| ***SLC17A6*** | 2930.37 | -0.01 | 0.11 | 0.91 | 0.96 | -0.00 | 0.04 |
| ***SLIT1*** | 12650.29 | -0.02 | 0.17 | 0.91 | 0.96 | -0.00 | 0.04 |
| ***ADCYAP1R1*** | 710.41 | -0.02 | 0.18 | 0.92 | 0.96 | -0.00 | 0.04 |
| ***ECEL1*** | 1127.51 | 0.02 | 0.21 | 0.93 | 0.97 | 0.00 | 0.04 |
| ***CALCR*** | 696.66 | 0.01 | 0.11 | 0.95 | 0.97 | 0.00 | 0.04 |
| ***PMCH*** | 59.33 | -0.03 | 0.45 | 0.95 | 0.97 | -0.00 | 0.05 |
| ***HMX3*** | 1714.50 | -0.01 | 0.19 | 0.97 | 0.98 | -0.00 | 0.04 |
| ***EMX2*** | 917.69 | -0.01 | 0.26 | 0.98 | 0.98 | -0.00 | 0.05 |
| ***CPNE2*** | 1755.28 | -0.00 | 0.11 | 0.98 | 0.98 | -0.00 | 0.04 |

**Table S2:** Genome-wide differential gene expression in medial hypothalamic punch RNA-seq from heat call-programmed zebra finches.

| Gene | Base Mean | Log2FoldChange | lfcSE | Stat | pvalue | padj |
|---|---|---|---|---|---|---|
| ***LOC115493855*** | 102.92 | -1.97 | 0.37 | 28.24 | 1.07E-07 | 1.93E-03 |
| ***SLC25A4*** | 2833.02 | -1.29 | 0.26 | 25.06 | 5.55E-07 | 3.37E-03 |
| ***TNNC2*** | 8193.28 | -1.83 | 0.36 | 25.04 | 5.61E-07 | 3.37E-03 |
| ***DHRS7C*** | 142.13 | -1.44 | 0.32 | 20.32 | 6.56E-06 | 2.36E-02 |
| ***KLHL31*** | 465.20 | -1.79 | 0.40 | 20.44 | 6.15E-06 | 2.36E-02 |

| | | | | | | |
|---|---|---|---|---|---|---|
| ***ACTA1*** | 25842.32 | -1.88 | 0.43 | 18.79 | 1.46E-05 | 2.45E-02 |
| ***FAM180B*** | 5.82 | -3.19 | 0.71 | 19.12 | 1.23E-05 | 2.45E-02 |
| ***FBXL22*** | 223.41 | -1.39 | 0.33 | 18.58 | 1.63E-05 | 2.45E-02 |
| ***KLHL40*** | 498.57 | -1.56 | 0.36 | 19.30 | 1.12E-05 | 2.45E-02 |
| ***LDB3*** | 2132.07 | -1.59 | 0.36 | 19.18 | 1.19E-05 | 2.45E-02 |
| ***LOC115495486*** | 137.86 | -1.97 | 0.48 | 19.11 | 1.23E-05 | 2.45E-02 |
| ***MUSTN1*** | 803.68 | -1.62 | 0.37 | 18.68 | 1.55E-05 | 2.45E-02 |
| ***MYOM1*** | 1367.09 | -1.50 | 0.36 | 18.27 | 1.92E-05 | 2.6E-02 |
| ***PACSIN3*** | 1502.59 | -1.13 | 0.27 | 18.17 | 2.02E-05 | 2.6E-02 |
| ***CMYA5*** | 390.52 | -1.52 | 0.37 | 18.02 | 2.18E-05 | 2.62E-02 |
| ***MYOT*** | 1360.14 | -1.54 | 0.36 | 17.81 | 2.45E-05 | 2.75E-02 |
| ***FSD2*** | 154.26 | -1.35 | 0.33 | 17.51 | 2.86E-05 | 2.83E-02 |
| ***LOC101233779*** | 123.12 | -1.34 | 0.32 | 17.43 | 2.99E-05 | 2.83E-02 |
| ***NEB*** | 2167.99 | -1.77 | 0.43 | 17.47 | 2.92E-05 | 2.83E-02 |
| ***CLIC5*** | 70.57 | -1.16 | 0.29 | 17.25 | 3.28E-05 | 2.96E-02 |
| ***OBSCN*** | 878.09 | -1.36 | 0.34 | 17.02 | 3.7E-05 | 3.17E-02 |
| ***ATP1B4*** | 347.24 | -1.29 | 0.32 | 16.90 | 3.95E-05 | 3.23E-02 |
| ***CASQ2*** | 2358.59 | -1.73 | 0.43 | 16.63 | 4.55E-05 | 3.5E-02 |
| ***LOC105758604*** | 7951.80 | 0.20 | 0.05 | 16.43 | 5.06E-05 | 3.5E-02 |
| ***LOC115493912*** | 38.21 | -1.54 | 0.41 | 16.52 | 4.81E-05 | 3.5E-02 |
| ***LOC121468823*** | 15.09 | -1.80 | 0.44 | 16.46 | 4.96E-05 | 3.5E-02 |
| ***TNNT3*** | 23912.25 | -1.43 | 0.36 | 15.88 | 6.73E-05 | 4.37E-02 |
| ***TRIM55*** | 170.78 | -1.48 | 0.39 | 15.87 | 6.8E-05 | 4.37E-02 |
| ***UNC45B*** | 515.04 | -1.32 | 0.34 | 15.76 | 7.2E-05 | 4.39E-02 |
| ***ZNF106*** | 2405.54 | -0.79 | 0.21 | 15.73 | 7.32E-05 | 4.39E-02 |
| ***MYLK2*** | 213.82 | -1.43 | 0.39 | 15.53 | 8.1E-05 | 4.45E-02 |
| ***MYOZ1*** | 442.64 | -1.95 | 0.50 | 15.52 | 8.16E-05 | 4.45E-02 |
| ***TNNI2*** | 4836.21 | -1.69 | 0.43 | 15.58 | 7.9E-05 | 4.45E-02 |
| ***ASB12*** | 238.01 | -1.38 | 0.36 | 15.33 | 9.04E-05 | 4.65E-02 |
| ***PPP1R3A*** | 179.42 | -1.40 | 0.37 | 15.36 | 8.89E-05 | 4.65E-02 |
| ***LOC100218875*** | 2095.71 | -1.81 | 0.49 | 15.18 | 9.79E-05 | 4.67E-02 |
| ***SLC22A16*** | 141.68 | -1.25 | 0.34 | 15.19 | 9.7E-05 | 4.67E-02 |

| | | | | | | |
|---|---|---|---|---|---|---|
| ***XIRP1*** | 591.38 | -1.56 | 0.41 | 15.11 | 1.01E-04 | 4.67E-02 |
| ***YIPF7*** | 54.77 | -0.85 | 0.22 | 15.13 | 1E-04 | 4.67E-02 |
| ***FHL1*** | 2000.44 | -1.78 | 0.45 | 15.06 | 1.04E-04 | 4.69E-02 |
| ***SRL*** | 2776.20 | -1.38 | 0.37 | 14.98 | 1.09E-04 | 4.78E-02 |
| ***CACNG1*** | 755.42 | -1.35 | 0.35 | 14.84 | 1.17E-04 | 4.81E-02 |
| ***EMILIN3*** | 680.55 | -0.50 | 0.13 | 14.71 | 1.26E-04 | 4.81E-02 |
| ***LOC100228485*** | 1664.12 | -2.32 | 0.56 | 14.87 | 1.15E-04 | 4.81E-02 |
| ***LOC100230937*** | 32.60 | -1.45 | 0.39 | 14.77 | 1.21E-04 | 4.81E-02 |
| ***LOC105760992*** | 29574.71 | -1.93 | 0.50 | 14.71 | 1.25E-04 | 4.81E-02 |
| ***TPM2*** | 16134.92 | -1.06 | 0.28 | 14.82 | 1.19E-04 | 4.81E-02 |
| ***LOC115495214*** | 50.13 | -1.77 | 0.47 | 14.64 | 1.3E-04 | 4.88E-02 |
| ***DES*** | 2910.36 | -1.43 | 0.36 | 14.57 | 1.35E-04 | 4.95E-02 |

**Table S3:** Gene Ontology Biological Process enrichment results for modules correlated with heat call playback. Green, brown, and red module genes were analyzed using ShinyGO (v0.85, zebra finch) using hypergeometric tests with Benjamini–Hochberg FDR correction. Number of genes, fold enrichment, and a list of enriched genes are provided for each significant GO term (the brown module did not return significant GO term).

| Pathway | Enrichment FDR | Module | Number of Genes | Pathway Genes | Fold Enrichment | Genes |
|---|---|---|---|---|---|---|
| **Path:hsa04820 Cytoskeleton in muscle cells** | 2.64E-33 | Green | 48 | 232 | 10.21 | *PDLIM5, LDB3, MYOM3, COL6A2, DES, SYNPO2, FHL1, ATP1B4, ANKRD2, AMPD1, PDLIM3, LMNA, LMOD2, MYBPC1, MYBPC3, MYL1, MYL2, MYL3, NEB, MYOZ2, LMOD3, ACTA1, MYOZ1, SGCD, SGCG, SNTB1, ACTC1, TMOD1, TNNC2, TNNC1, TNNI1, TNNI2, TNNT2, TNNT3, TPM1, TPM2, TPM4, VIM, CSRP3, SSPN, CAPN3, MYPN, TRIM55, MYOM1, ACTN2, PDLIM1, MYOM2, MYOT* |
| **Path:hsa04260 Cardiac muscle contraction** | 5.74E-06 | Green | 11 | 87 | 7.83 | *ATP1B4, MYL2, MYL3, ACTC1, TNNC1, TNNT2, TPM1, TPM2, TPM4, CACNG1, CASQ2* |
| **Path:hsa05410 Hypertrophic cardiomyopathy** | 8.89E-09 | Green | 17 | 99 | 7.42 | *DES, LMNA, MYBPC3, MYL2, MYL3, SGCD, SGCG, SNTB1, ACTC1, TNNC1, TNNT2, TPM1, TPM2, TPM4, CACNG1, SSPN, CAV3* |
| **Path:hsa05414 Dilated cardiomyopathy** | 1.10E-08 | Green | 17 | 105 | 7.15 | *DES, LMNA, MYBPC3, MYL2, MYL3, SGCD, SGCG, SNTB1, ACTC1, TNNC1, TNNT2, TPM1, TPM2, TPM4, CACNG1, SSPN, CAV3* |
| **Path:hsa05416 Viral myocarditis** | 1.23E-02 | Green | 6 | 69 | 5.98 | *SGCD, SGCG, SNTB1, SSPN, CAV1, CAV3* |

| | | | | | | |
|---|---|---|---|---|---|---|
| **Path:hsa04814 Motor proteins** | 1.73E-05 | Green | 16 | 194 | 4.58 | *DYNC1I1, MYL1, MYL2, MYL3, MYO5C, ACTA1, ACTC1, TNNC2, TNNC1, TNNI1, TNNI2, TNNT2, TNNT3, TPM1, TPM2, TPM4* |
| **Path:hsa05412 Arrhythmogenic right ventricular cardiomyopathy** | 5.37E-03 | Green | 9 | 86 | 4.49 | *DES, LMNA, SGCD, SGCG, SNTB1, CACNG1, SSPN, CAV3, ACTN2* |
| **Path:hsa04261 Adrenergic signaling in cardiomyocytes** | 5.23E-04 | Green | 13 | 153 | 4.16 | *BVES, AGTR1, ATP1B4, MYL2, MYL3, POPDC2, ACTC1, TNNC1, TNNT2, TPM1, TPM2, TPM4, CACNG1* |
| **GO:0006936 Muscle contraction** | 4.25E-02 | Red | 19 | 360 | 2.91 | *CHRM2, SRF, ATP2A1, MAP2K3, ARG2, CACNA1S, CHRND, ATP1B1, TRIM63, ATP1A1, ABAT, ANXA6, SNTA1, FGF12, NMUR2, CRYAB, GAMT, MYBPH, CHRNG* |
| **GO:0003012 Muscle system process** | 2.35E-02 | Red | 23 | 450 | 2.75 | *CHRM2, SRF, ATP2A1, MAP2K3, ACACB, ARG2, CACNA1S, MYOG, HEY2, CHRND, ATP1B1, TRIM63, ATP1A1, ABAT, MYMK, ANXA6, SNTA1, FGF12, NMUR2, CRYAB, GAMT, MYBPH, CHRNG* |
| **GO:0061061 Muscle structure development** | 1.69E-02 | Red | 33 | 721 | 2.35 | *MYF5, MYOG, PDLIM4, MYBPH, ALPK3, DOCK1, PDGFB, MYMK, SOD2, EHD2, YBX3, CACNA1S, CRYAB, SRF, MEF2D, TCF21, WFIKKN1, FLNC, HOXD9, AKIRIN2, HEY2, SYPL2, UCHL1, FBXO40, HSPB2, TMEM119, PPP2R3A, CHRND, BOC, OBSL1, CHODL, ITGB1BP2, FHL3* |

**Table S4:** Significant isoform switches (FDR < 0.05, differential isoform fraction ≥ 0.05) in medial hypothalamic RNA-seq from embryos exposed to chronic heat call versus control playback.

| Isoform ID | Gene Symbol | Reference Condition | Treatment Condition | Isoform Fraction in Control | Isoform Fraction in Heat Call | Change in isoform fraction (ΔIF) | q-value |
|---|---|---|---|---|---|---|---|
| **XM_041721404.1** | *TLK2* | cont | treat | 0.06 | 0 | -0.06 | 9.4E-07 |
| **XM_041714671.1** | *LOC100228270* | cont | treat | 0.33 | 0.07 | -0.26 | 8.6E-03 |
| **NM_001136481.2** | *TPM1* | cont | treat | 0.66 | 0.43 | -0.23 | 8.6E-03 |
| **XM_030265025.3** | *TCAIM* | cont | treat | 0.17 | 0.02 | -0.15 | 1.2E-02 |
| **XM_030285413.3** | *EIF4ENIF1* | cont | treat | 0.21 | 0.38 | 0.17 | 1.6E-02 |
| **XM_041721341.1** | *CELSR2* | cont | treat | 0.78 | 0.58 | -0.2 | 2.4E-02 |
| **XM_030269705.3** | *GPCPD1* | cont | treat | 0.44 | 0.64 | 0.21 | 2.4E-02 |
| **XM_041721627.1** | *MPV17L2* | cont | treat | 0.03 | 0.39 | 0.36 | 2.4E-02 |
| **XM_041721628.1** | *MPV17L2* | cont | treat | 0.97 | 0.61 | -0.36 | 2.4E-02 |
| **XM_030272879.3** | *RLIM* | cont | treat | 0.71 | 0.88 | 0.17 | 2.4E-02 |
| **XM_030272880.3** | *RLIM* | cont | treat | 0.29 | 0.1 | -0.18 | 2.4E-02 |
| **XM_002191938.4** | *TSPEAR* | cont | treat | 0.84 | 0.94 | 0.1 | 2.4E-02 |

| | | | | | | | |
|---|---|---|---|---|---|---|---|
| **XM_041717575.1** | *CARF* | cont | treat | 0.08 | 0.02 | -0.07 | 3E-02 |
| **XM_030280110.3** | *ARHGEF7* | cont | treat | 0.16 | 0.04 | -0.12 | 3.3E-02 |
| **XM_030286601.3** | *RGS3* | cont | treat | 0.08 | 0.16 | 0.07 | 3.3E-02 |
| **XM_030280436.3** | *TSPEAR* | cont | treat | 0.16 | 0.06 | -0.1 | 3.3E-02 |
| **XM_030266254.3** | *LOC121468029* | cont | treat | 0.27 | 0.14 | -0.13 | 3.6E-02 |
| **XM_030271401.3** | *CBR4* | cont | treat | 0.07 | 0.01 | -0.06 | 3.9E-02 |
| **XM_030263200.3** | *CHPT1* | cont | treat | 0.11 | 0.05 | -0.06 | 3.9E-02 |
| **XM_030284686.3** | *GSG1L* | cont | treat | 0.16 | 0.05 | -0.11 | 3.9E-02 |
| **XM_030267870.3** | *PTPRK* | cont | treat | 0.34 | 0.55 | 0.21 | 3.9E-02 |
| **XM_012573306.4** | *TBC1D9* | cont | treat | 0.28 | 0.16 | -0.12 | 3.9E-02 |
| **XM_030271911.3** | *TBC1D9* | cont | treat | 0.72 | 0.84 | 0.12 | 3.9E-02 |
| **XM_002200039.6** | *ARFGAP2* | cont | treat | 0.91 | 0.98 | 0.07 | 3.9E-02 |
| **XM_030283390.3** | *DENND6A* | cont | treat | 0 | 0.24 | 0.24 | 3.9E-02 |
| **XM_030275365.3** | *LOC100222881* | cont | treat | 0.11 | 0.4 | 0.29 | 3.9E-02 |
| **XM_030275366.3** | *LOC100222881* | cont | treat | 0.89 | 0.6 | -0.29 | 3.9E-02 |
| **XM_030274227.3** | *ARFGAP2* | cont | treat | 0.09 | 0.02 | -0.07 | 4E-02 |
| **XM_002189844.6** | *DHTKD1* | cont | treat | 0.9 | 0.73 | -0.17 | 4.4E-02 |
| **XM_032751614.2** | *RAB37* | cont | treat | 0.42 | 0.08 | -0.34 | 4.4E-02 |
| **XM_041717123.1** | *LOC121470179* | cont | treat | 0.24 | 0.01 | -0.23 | 4.7E-02 |
| **XM_030290503.3** | *DHTKD1* | cont | treat | 0.1 | 0.27 | 0.17 | 4.9E-02 |
| **XM_030288176.3** | *ANKFY1* | cont | treat | 0.45 | 0.25 | -0.2 | 4.9E-02 |
| **XM_030288177.3** | *ANKFY1* | cont | treat | 0.55 | 0.75 | 0.2 | 4.9E-02 |

**Table S5:** Alternative splicing event enrichment among significant isoform switches in medial hypothalamic RNA-seq from heat call and control embryos.

| Splicing Type | Number of Gains in Heat Call | Number of Losses in Heat Call | Total Events | Proportion of Gains | P-value | q-value |
|---|---|---|---|---|---|---|
| **ES (Exon Skipping)** | 4 | 13 | 17 | 0.24 | 0.049 | 0.34 |
| **A3SS (Alt 3' Splice Site)** | 5 | 1 | 6 | 0.83 | 0.219 | 0.77 |
| **A5SS (Alt 5' Splice Site)** | 0 | 1 | 1 | 0 | 1.000 | 1.00 |

| | | | | | | |
|---|---|---|---|---|---|---|
| **ATSS (Alt Transcription Start)** | 6 | 3 | 9 | 0.67 | 0.508 | 1.00 |
| **ATTS (Alt Transcription Term)** | 3 | 2 | 5 | 0.6 | 1.000 | 1.00 |
| **IR (Intron Retention)** | 0 | 1 | 1 | 0 | 1 | 1 |
| **MES (Mutually Exclusive Exons)** | 3 | 2 | 5 | 0.6 | 1.000 | 1.00 |

**Table S6:** CIBERSORTx absolute-mode cell-type scores and deconvolution diagnostics for bulk hypothalamic RNA-seq samples.

| Mixture | astrocyte | ependymal cell | oligodendrocyte | neuron | endothelial cell | microglial cell | fibroblast | mural cell | P-value | Correlation | RMSE | Absolute score (sig.score) |
|---|---|---|---|---|---|---|---|---|---|---|---|---|
| **S08_Treat_F** | 0.180 | 0.002 | 0.692 | 0.746 | 0.013 | 0.098 | 0.000 | 0.007 | 0.000 | 0.977 | 0.700 | 1.738 |
| **S10_Treat_F** | 0.193 | 0.052 | 0.673 | 0.658 | 0.003 | 0.109 | 0.000 | 0.008 | 0.000 | 0.894 | 0.749 | 1.698 |
| **S12_Cont_M** | 0.238 | 0.005 | 0.683 | 0.559 | 0.001 | 0.136 | 0.000 | 0.071 | 0.000 | 0.914 | 0.750 | 1.693 |
| **S13_Cont_M** | 0.224 | 0.017 | 0.665 | 0.650 | 0.004 | 0.122 | 0.000 | 0.022 | 0.000 | 0.962 | 0.721 | 1.704 |
| **S14_Cont_M** | 0.189 | 0.028 | 0.691 | 0.628 | 0.006 | 0.125 | 0.000 | 0.013 | 0.000 | 0.974 | 0.719 | 1.680 |
| **S15_Treat_F** | 0.173 | 0.039 | 0.655 | 0.805 | 0.009 | 0.104 | 0.000 | 0.008 | 0.000 | 0.903 | 0.732 | 1.792 |
| **S16_Treat_M** | 0.192 | 0.017 | 0.682 | 0.762 | 0.005 | 0.118 | 0.000 | 0.003 | 0.000 | 0.968 | 0.707 | 1.780 |
| **S20_Treat_F** | 0.207 | 0.007 | 0.730 | 0.551 | 0.007 | 0.114 | 0.000 | 0.050 | 0.000 | 0.898 | 0.754 | 1.665 |
| **S21_Cont_M** | 0.218 | 0.030 | 0.706 | 0.598 | 0.012 | 0.127 | 0.000 | 0.024 | 0.000 | 0.956 | 0.731 | 1.715 |
| **S22_Treat_F** | 0.216 | 0.000 | 0.669 | 0.654 | 0.012 | 0.121 | 0.000 | 0.030 | 0.000 | 0.981 | 0.711 | 1.703 |
| **S24_Treat_M** | 0.192 | 0.001 | 0.659 | 0.679 | 0.016 | 0.134 | 0.000 | 0.017 | 0.000 | 0.975 | 0.711 | 1.698 |
| **S25_Cont_M** | 0.200 | 0.012 | 0.684 | 0.661 | 0.011 | 0.097 | 0.000 | 0.015 | 0.000 | 0.972 | 0.711 | 1.680 |
| **S26_Treat_F** | 0.292 | 0.004 | 0.734 | 0.541 | 0.017 | 0.126 | 0.005 | 0.025 | 0.000 | 0.953 | 0.735 | 1.744 |
| **S27_Cont_F** | 0.215 | 0.057 | 0.665 | 0.565 | 0.004 | 0.119 | 0.000 | 0.052 | 0.000 | 0.961 | 0.734 | 1.678 |
| **S28_Treat_F** | 0.197 | 0.011 | 0.695 | 0.672 | 0.007 | 0.099 | 0.000 | 0.015 | 0.000 | 0.969 | 0.712 | 1.695 |
| **S30_Cont_M** | 0.260 | 0.016 | 0.675 | 0.481 | 0.018 | 0.142 | 0.000 | 0.072 | 0.000 | 0.939 | 0.749 | 1.663 |
| **S31_Treat_M** | 0.204 | 0.011 | 0.706 | 0.613 | 0.010 | 0.114 | 0.000 | 0.031 | 0.000 | 0.974 | 0.719 | 1.689 |
| **S32_Cont_F** | 0.253 | 0.014 | 0.742 | 0.517 | 0.011 | 0.144 | 0.012 | 0.038 | 0.000 | 0.955 | 0.735 | 1.731 |
| **S33_Cont_M** | 0.221 | 0.004 | 0.671 | 0.604 | 0.009 | 0.129 | 0.000 | 0.041 | 0.000 | 0.977 | 0.719 | 1.678 |
